# Revealing the Widespread Bias of Extinction Risk in the Antarctic and the Southern Ocean

**DOI:** 10.64898/2026.04.23.720284

**Authors:** Madison G. Farrant, W. P. Amy Liu, Melodie A. McGeoch

## Abstract

Accelerating environmental change in the Antarctic and Southern Ocean (ASO) necessitates robust extinction risk assessments to inform conservation priorities and track progress towards global biodiversity targets. Nevertheless, no systematic, region-wide baseline of extinction risk currently exists for tracking ASO biodiversity responses to ongoing change, a significant barrier to global biodiversity monitoring. Here, we present the first comprehensive synthesis of extinction risk knowledge spanning plants, animals, and fungi across the ASO, examining biases in current assessments, the distribution of Threatened species and their associated threats. In the absence of a complete regional species checklist, species were compiled from >6,800,000 occurrences and existing checklists, yielding 5,403 assessments representing 2,806 species using a data-inclusive workflow that increased available assessments by over three-fold. Assessments are heavily biased towards vertebrates (56% assessed), while invertebrates, despite their ecological prevalence, are markedly underrepresented (4% assessed). Among vertebrates, mammals have the highest proportion of Threatened species (35%), while ASO birds are disproportionately Threatened (27%) compared to the global average (12%) with the greatest threat for ASO species being ‘Biological Resource Use’. Despite more Threatened species in the sub-Antarctic islands and the Antarctic Peninsula, relative to assessment effort, these regions had fewer Threatened species than expected, indicating these areas may function as refugia. These pronounced assessment biases highlight the need for more balanced, representative, and data-inclusive extinction risk assessments to be able to effectively detect conservation status change. This work represents an important step in ensuring ASO representation in global biodiversity monitoring frameworks strengthening the capacity of these frameworks to detect, attribute, and respond to future biodiversity changes.

## Introduction

Biodiversity decline in the Antarctic and Southern Ocean (ASO) is an increasingly urgent concern. Extinction risk assessments are fundamental tools for monitoring species trends (Hoffmann et al., 2010; Sayer et al., 2025), identifying threatened taxa (Possingham et al., 2002), and guiding conservation resource allocation (Guénard et al., 2025; Rodrigues et al., 2006), providing critical means of detecting biological responses pressures across regions. Assessments also determine the urgency with which conservation actions are implemented (Bachman et al., 2019). Species classified as ‘Threatened’ are often recognised in policy (Possingham et al., 2002; Rondinini et al., 2014) and public awareness gained from a Threatened classification has been linked to reduced extinction rates (Butchart et al., 2006; Edgar, 2025). Robust extinction risk evaluation is therefore fundamental to both understanding and responding to accelerating global change.

The value of extinction risk assessments depends on the breadth and completeness of their coverage. Despite this, substantial taxonomic, geographic, and ecological biases remain (Bachman et al., 2019; Rodrigues et al., 2006), with ASO thought to be underrepresented (Hughes et al., 2024; Lee et al., 2022) and often excluded from ‘global’ analyses of extinction risk and Threatened species trends (Gallagher et al., 2020; Hoffmann et al., 2010; Howard et al., 2020; Sayer et al., 2025). This omission is striking considering the ASO is a region characterised by exceptional levels of endemism, ecological distinctiveness, and productivity (Constable et al., 2003; Convey & Peck, 2019; Dehling & Chown, 2025), alongside its well-documented sensitivity to change (Brasier et al., 2021; Convey & Peck, 2019). Furthermore, the ASO also includes some of the most rapidly changing climates globally (Beale et al., 2026; Gorodetskaya et al., 2023; Morley et al., 2020), in addition to increased pressure from human activity (Brooks et al., 2019; Leihy et al., 2020), and invasive species (Lee et al., 2022; Leihy et al., 2023) which collectively intensifies extinction risk. Under high emissions scenarios, species protection has been identified as one of the most beneficial strategies for Antarctic conservation (Lee et al., 2022). Yet, without an ASO region-wide analysis, the true scale of species risk and the effectiveness of protection strategies remains unclear.

Addressing this information shortfall is critical not only for improving scientific understanding and regional species management, but for fulfilling both local and global conservation policy frameworks. Regionally, revisions to Annex II of the Antarctic Treaty System’s Committee for Environmental Protection (CEP) specify the use of International Union for the Conservation of Nature (IUCN) extinction risk information when designating Antarctic Specially Protected Species (SPS) (CEP, 2024). Globally, the Kunming-Montreal Global Biodiversity Framework (GBF) was adopted in 2022, sets action-oriented targets directly relevant to the ASO, including Target 4 aims to halt human-induced extinctions and reduce the extinction risk of Threatened species by 2030, while Goal A seeks to reduce extinction rates of all species by 2050 (Convention on Biological Diversity, 2022). In recognition of this, the 46^th^ Antarctic Treaty Consultative Meeting (ATCM) called for a better understanding of ASO species extinction risk (CEP, 2024). Nonetheless, no coordinated strategy across Antarctic governance parties has been implemented to advance this knowledge and provide a framework for assessing the progress of the GBF within the ASO.

An ASO-specific assessment of extinction risk knowledge and Threatened species will support decision-makers and deliver evidence for prioritising and implementing effective threat management and conservation responses (Chown et al., 2022), thereby enhancing progress towards conservation frameworks including the global GBF and local designation of Antarctic SPS. Here, we conduct the first ASO-wide assessment of current extinction risk, establishing a baseline essential for monitoring future biodiversity trends. Specifically, we: (1) identify current data gaps and biases in ASO extinction risk coverage; (2) examine the status of Threatened species within the ASO and evaluate associated threats; and (3) determine the extent to which Threatened species are protected by existing area-based conservation measures. Together, these results identify key management actions to strengthen the capacity of GBF’s to address biodiversity change across the region.

## Methods

### Study Area

The delineation of the Antarctic and Southern Ocean (ASO) region follows that adopted in the Marine Ecosystem Assessment for the Southern Ocean (MEASO) (Constable et al., 2023). Developed in 2018, MEASO represented the first quantitative circumpolar assessment of the status and trends of the Southern Ocean ecosystem which includes the Antarctic continent, the Southern Ocean, and several sub-Antarctic islands, including the Kerguelen Islands, Heard Island, and Campbell Island/Motu Ihupuku. The area included in the MEASO region was designed to be representative of the environmental and ecological heterogeneity characterising Antarctica, the Southern Ocean, and the sub-Antarctic (Constable et al., 2023).

### Data Acquisition

We collated and examined assessments for species in the kingdoms Animalia, Plantae, and Fungi using an integrated approach that included all extinction risk knowledge available to decision makers. Three primary datasets were compiled to analyse extinction risk assessments for both marine, terrestrial, and inland-water ASO species (Figure 1). All data acquisition and analyses were conducted using RStudio version 2024.04.0+735 (R Core Team, 2022). Detailed information on data downloads, record inclusion specifications, taxonomic harmonisation, and dataset cleaning for all three datasets is provided in Appendix S1. Datasets and R code for data acquisition and analyses are available through FigShare (https://doi.org/10.6084/m9.figshare.31175305) (Farrant et al., 2026) in accordance with FAIR (Findable, Accessible, Interoperable, and Reusable) data principles (Wilkinson et al., 2016).

**Figure 1:**
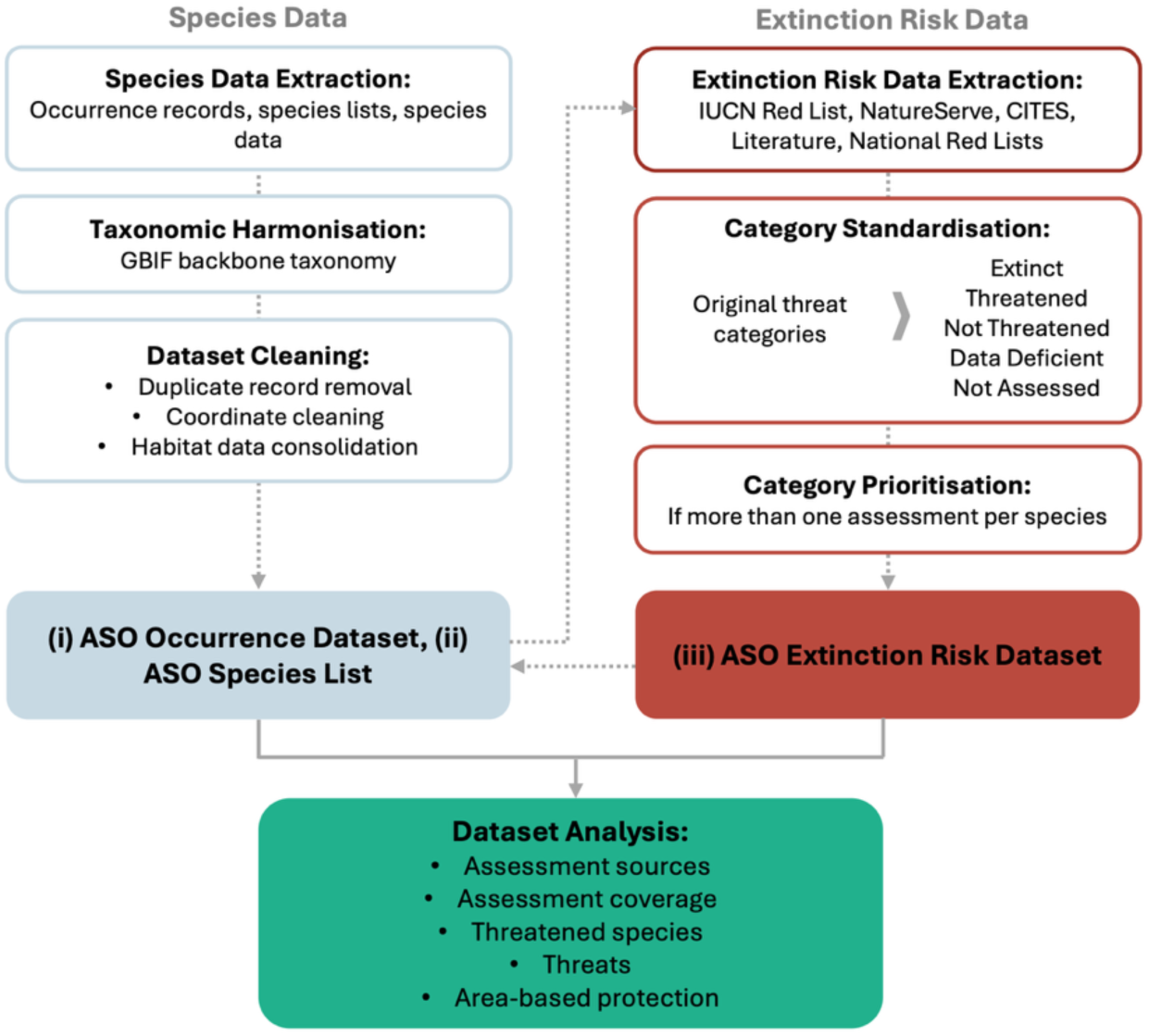
Data acquisition and integrative extinction risk workflow used in dataset compilation and analysis. Datasets produced are the (i) Antarctic and Southern Ocean (ASO) Occurrence Dataset, (ii) the ASO Species List, and (iii) the Extinction Risk Dataset.

The first dataset, the ‘Occurrence Dataset’, comprises georeferenced occurrence records for extant species within the ASO region, downloaded from four open-access databases (Table 1). Records were cleaned for duplicates, potential coordinate inaccuracies, and taxonomically harmonised using Global Biodiversity Information Facility (GBIF) Backbone Taxonomy (GBIF Secretariat, 2023). The final Occurrence Dataset (Occurrence_dataset.csv) contained 6,819,007 species-level occurrences from the ASO region representing an inventory of marine, terrestrial, and inland-water dwelling species, with observation dates spanning 1697 to 2025. Each record includes information on species taxonomy, occurrence status, observation date, habitat, extinction risk, and data source(s). The Occurrence Dataset served as the foundation for identifying species present within the ASO region, in the absence of a comprehensive checklist for the MEASO region. It was also used for assessing spatial patterns in species assessments and to map the spatial distribution of Threatened taxa and protected areas within the ASO.

**Table 1:**
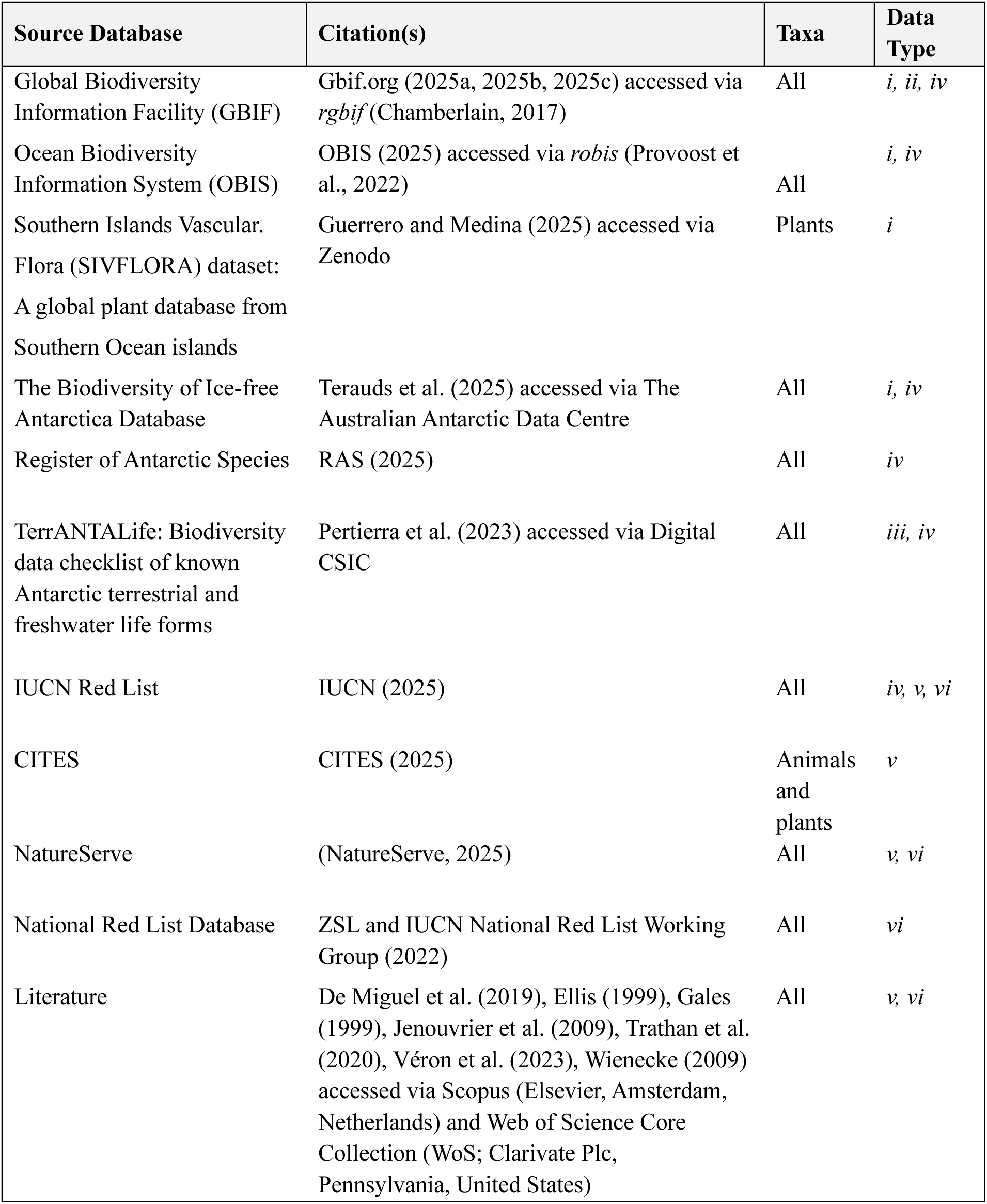
Sources used to compile the Occurrence Dataset, the Antarctic and Southern Ocean (ASO) Species List, and the Extinction Risk Datasets.

| Source Database | Citation(s) | Taxa | Data Type |
| --- | --- | --- | --- |
| Global Biodiversity Information Facility (GBIF) | Gbif.org (2025a, 2025b, 2025c) accessed via *rgbif* (Chamberlain, 2017) | All | *i, ii, iv* |
| Ocean Biodiversity Information System (OBIS) | OBIS (2025) accessed via *robis* (Provoost et al., 2022) | All | *i, iv* |
| Southern Islands Vascular Flora (SIVFLORA) dataset: A global plant database from Southern Ocean islands | Guerrero and Medina (2025) accessed via Zenodo | Plants | *i* |
| The Biodiversity of Ice-free Antarctica Database | Terauds et al. (2025) accessed via The Australian Antarctic Data Centre | All | *i, iv* |
| Register of Antarctic Species | RAS (2025) | All | *iv* |
| TerrANTALife: Biodiversity data checklist of known Antarctic terrestrial and freshwater life forms | Pertierra et al. (2023) accessed via Digital CSIC | All | *iii, iv* |
| IUCN Red List | IUCN (2025) | All | *iv, v, vi* |
| CITES | CITES (2025) | Animals and plants | *v* |
| NatureServe | (NatureServe, 2025) | All | *v, vi* |
| National Red List Database | ZSL and IUCN National Red List Working Group (2022) | All | *vi* |
| Literature | De Miguel et al. (2019), Ellis (1999), Gales (1999), Jenouvrier et al. (2009), Trathan et al. (2020), Véron et al. (2023), Wienecke (2009) accessed via Scopus (Elsevier, Amsterdam, Netherlands) and Web of Science Core Collection (WoS; Clarivate Plc, Pennsylvania, United States) | All | *v, vi* |

The second dataset created, the ‘ASO Species List’, provides a taxonomically harmonised list of ASO species sourced from the Occurrence Dataset, combined with existing species checklists (Table 1), including the ‘TerrANTALife’ biodiversity checklist of known Antarctic terrestrial and freshwater life forms (Pertierra et al., 2023, 2024). The final ASO Species List (ASO_species_list.csv) contains 20,764 species spanning 42 phyla. As with the Occurrence Dataset, the ASO Species List also compiles information on species taxonomy, habitat, extinction risk, and data source(s) and was intended act as a searchable list of species for the extraction of extinction risk assessments and their related attributes. We acknowledge that, due to the nature of using occurrences to derive the species list, despite best data screening efforts, some inaccuracies will be inevitable. In addition, recently described species or taxonomic changes may not have been captured within this study, due to a lag between formal species description, taxonomic revision, and integration into the GBIF Backbone Taxonomy. Nevertheless, we believe the ASO Species List presented here represents the best estimate of the ASO species pool and encourage further research to refine and expand this list.

Extinction risk assessments for species listed in the ASO Species List were extracted from four databases and supplemented through literature searches (Table 1). These assessments were consolidated into a third dataset, the ‘Extinction Risk Dataset’, which enabled detailed evaluation of assessment sources and conservation status. To examine current progress towards GBF goals and targets, we incorporated all available extinction risk assessments for ASO species because biodiversity research remains comparatively limited (Koerich et al., 2023; Pertierra et al., 2025), maximising use of all available information for supporting conservation decision-making. This included assessments conducted at multiple scales, encompassing both population-level (national to sub-global scale assessments) and species-level assessments (global scale assessments) in addition to multiple assessment sources (Holz et al., 2022; Lončarević et al., 2024; Mounce et al., 2018). This included 61 different publications (Appendix S2) consolidated from the National Red List Database (ZSL and IUCN National Red List Working group, 2022). A data-inclusive approach is particularly valuable in regions that are data-limited, such as the ASO (Koerich et al., 2023), allowing a more nuanced evaluation of conservation status through revealing hidden declines which may be masked at the species level (Brito et al., 2010; Mounce et al., 2018).

To address variability in the lexicon used between assessment sources, assessment statuses were standardised into five conservation statuses: Extinct, Threatened, Not Threatened, Not Assessed, and Data Deficient (Appendix S3). The original criteria used by each source were reviewed to ensure that the standardised conservation status accurately reflected the intended published conservation status. For subsequent data analyses on Threatened species, in the instances where species had multiple conservation statuses, priority was given to the conservation status of assessments completed within the last 10 years, then was conservatively assigned to the most Threatened category (Appendix S4) (from most to least prioritised: Extinct, Threatened, Not Threatened, Data Deficient, then Not Assessed) for a conservative approach. The final Extinction Risk Dataset (Extinction_risk_dataset.csv) contains all compiled extinction risk assessments for species within the ASO Species List. In addition, information on species taxonomy, assessment source, assessment scale, publication, and both the original, standardised, and prioritised assessments were included where available for ease of analysis.

All vocabulary used in the creation of the Occurrence Dataset, the ASO Species List, and the Extinction Risk Dataset were aligned with Darwin Core bioinformatic standards where possible, to ensure consistency and interoperability (Wieczorek et al., 2012). Additional non-standard fields were used to capture information on species extinction risk assessments and conservation status (Appendix S5). The current versions of the three datasets represent available data up until March 2025. These datasets are intended to be updated as additional species lists, occurrences, and extinction risk assessments become available.

### Source Overlap and Assessment Agreement

Potential benefits of using an integrative and data-inclusive approach were evaluated by comparing sources of extinction risk assessments to determine the contribution of unique information. The number of unique species assessed by each source and the proportion of assessments representing different habitats and taxonomic groups were quantified at both the species and population levels. The degree of agreement among standardised conservation statuses was quantified using Cohen’s Kappa statistic (Cohen, 1960). Pairwise comparisons were conducted between the two largest sources of assessments, i.e. the IUCN Red List and NatureServe, at both species- and population-levels, with agreement matrices subsequently generated to identify discrepancies across corresponding conservation statuses (Appendix S8). Further methodological details for data analyses are provided in Appendix S1.

### Extinction Risk Assessment Coverage

Taxonomic, habitat, temporal, and geographic coverage of extinction risk assessments were examined to identify potential biases in assessment effort and gaps in conservation status knowledge. Analyses were conducted using integrated results for an overview of the ASO ecosystem. Taxonomic coverage was quantified by the proportion of species assessed per major taxonomic group and taxonomic sub-groups. Habitat coverage was examined by comparing the proportion of assessed species classified within terrestrial, marine, and inland-water habitats. It is important to examine habitat coverage as different habitats may face very distinct threats (Hogue & Breon, 2022). Temporal patterns in assessments were examined by quantifying the accumulation of assessments over time. Assessments extracted from NatureServe did not include the assessment dates, therefore, NatureServe assessments were excluded from temporal analysis. Geographic coverage was analysed to identify potential spatial biases in assessment density and effort. Using the Occurrence Dataset, the distribution of total number of assessed species at a 100 km resolution was examined to see assessment density across the ASO. Correlation analyses were performed using Spearman Rank Correlation to examine relationships between assessment density and species richness. Assessed species as a proportion of total species richness within each cell was reported as assessment effort.

### Threatened Species

Taxonomic and habitat coverage of threatened species was assessed by calculating the proportion of species within each standardised conservation status for different taxonomic groups and habitat/s. Relationships between assigned conservation status and taxonomic or habitat/s groups were tested using chi-squared analysis followed by examining standardised residuals. This allowed identification of which taxa or habitats were more or less likely to be assigned each conservation status than expected by chance.

The proportion of threatened species was calculated for each taxon using methods established by the IUCN guidelines for reporting on the proportion of Threatened species (Annex version 1.2) to account for the uncertainty of Data Deficient species (IUCN, 2022). The mid-point represents the most realistic estimate of Threatened species proportions by excluding Data Deficient and Extinct species from the denominator. We also report on the lower bound (no Data Deficient species considered Threatened) and the upper bound (all Data Deficient species considered Threatened) as the range (IUCN, 2022).

The geographic coverage of species classified as Threatened within the ASO region was examined to identify areas of potentially elevated threat or high conservation value. Using the Occurrence Dataset, the number of Threatened species was summed at a 100 km resolution. To further account for regional variation in species richness and assessment density, weighted species richness was calculated. Weighted species richness quantifies the relationship between the number of Threatened species relative to the total richness of Data Sufficient species (Data Sufficient species = Number of species assessed - Data Deficient species) (Hoffmann et al., 2010). Standardised residuals were then calculated for each cell. Analysis of these standardised residuals enables identification of areas with more threatened species than predicted based on species richness and assessment effort (Hoffmann et al., 2010).

### Threats to Species

For assessments sourced from the IUCN Red List, threat information is required for all species-level assessments classified within the Threatened categories (Critically Endangered, Endangered, and Vulnerable) as well as the Extinct categories (Extinct and Extinct in the Wild) (IUCN, 2013). Species with available threat data (*n* = 775) were extracted from the IUCN Red List (Table 1). The proportion of species affected by each conservation status was calculated across taxonomic groups. For comparison, the proportion of species affected by each threat driver was also calculated for each taxonomic group across all species assessed by the IUCN Red List globally (IUCN, 2025). Threat data is coded according to the IUCN threat classification scheme version 3.3 (IUCN, 2025).

### Protected Area Coverage of Threatened Species

The relationship between protected area coverage and conservation status was assessed by quantifying the extent of terrestrial and marine area-based protection afforded to species within the ASO. Terrestrial protected areas included 76 Antarctic Specially Protected Areas (ASPAs) and six Antarctic Specially Managed Areas (ASMAs) (Antarctic Treaty Secretariat, 2025). Marine protected areas included two Marine Protected Areas (MPAs) established by the Commission for the Conservation of Antarctic Marine Living Resources (CAMMLR). Additionally, there are five United Nations Educational, Scientific and Cultural Organization (UNESCO) World Heritage Sites located within the ASO which encompass both marine and terrestrial environments. A complete list of protected areas that are located, whether partially or wholly within the ASO is provided in Appendix S6. Polygons of protected areas were used to calculate the area of overlap with each species’ extent of occurrence (EOO), based on records in the Occurrence Dataset for species with at least three occurrences (*n* = 11,845 species). Differences in marine and terrestrial (including inland-water) species and protected area overlap (dependent variable) among conservation statuses (independent variable) were assessed using a Kruskal-Wallis non-parametric test (Kruskal & Wallis, 1952). Post-hoc pairwise comparisons were then performed using Dunn’s test (Dunn, 1964) with Benjamini-Hochberg correction.

## Results

### Assessment Sources and Agreement

Using an integrative approach that incorporated species assessments from different sources (Table 1) with both species- and population-level assessments, the number of assessments within the Antarctic and Southern Ocean (ASO) was more than three times greater than provided by any single source and level. At the species level, the IUCN Red List was the largest contributor of assessments (*n* = 1,748) followed by NatureServe (966). In contrast, at the population-level, this contribution was reversed with NatureServe contributing the most assessments (1,226) followed by the IUCN Red List (832), and national red lists (573).

Examining the contribution of unique information from each source demonstrated that different sources tended to focus on different species (Figure 2a). At the species level, the IUCN Red List had the highest number of unique species assessed, with 80% of species not assessed by any other source, followed by NatureServe (66%). At the population level, the IUCN Red List was the largest contributor of unique species with 55% of species not assessed by any other source followed by the NatureServe (51%) and national red list species (38%). CITES did not contribute any unique species assessments that were not already assessed by other sources. The overall contribution of literature to unique species assessments was low (26 species- and four population-level assessments). Across all sources, 1,089 species (64%) were assessed at both species- and population-levels.

**Figure 2:**
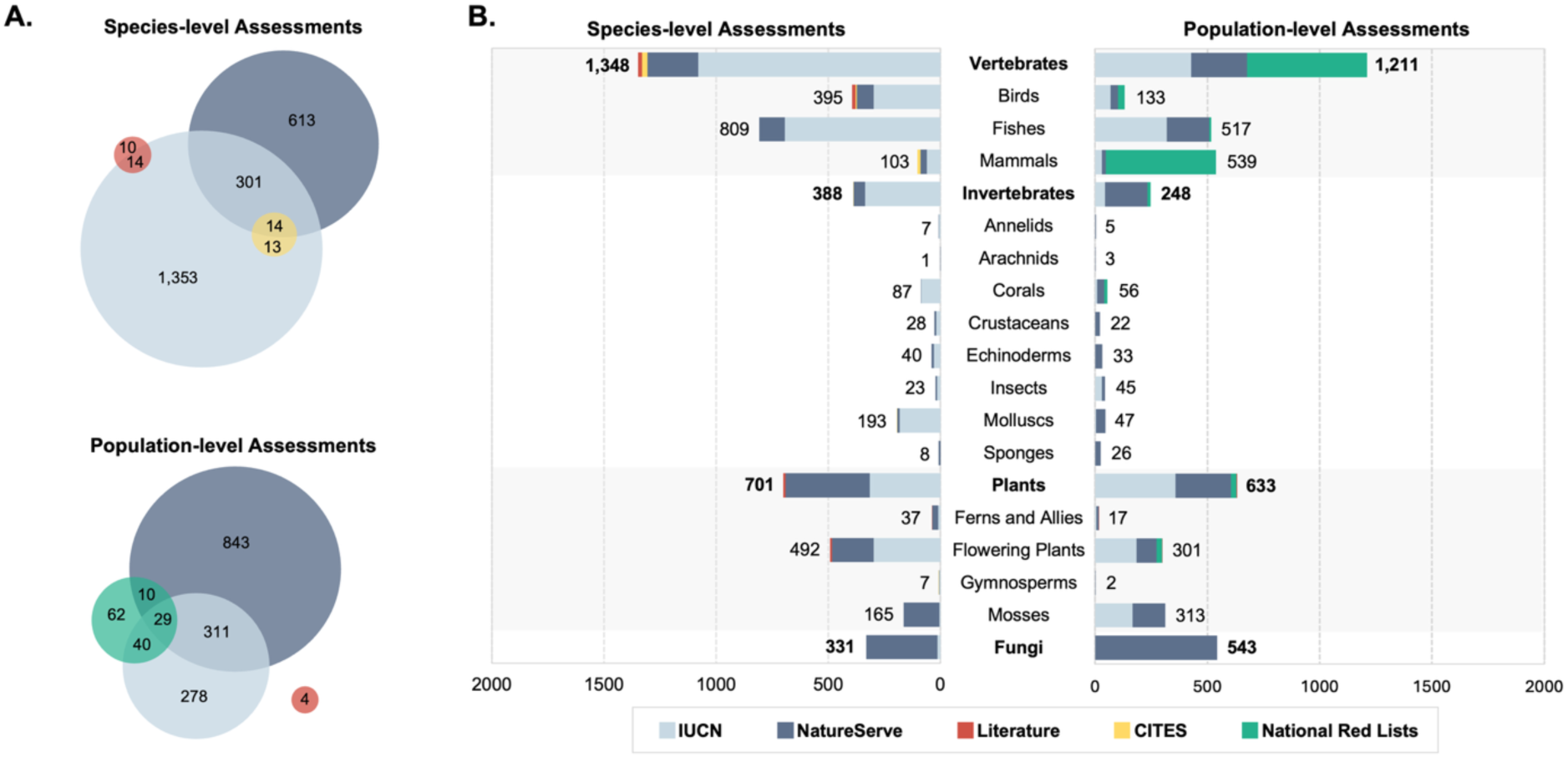
Sources of species- and population-level extinction risk assessments for Antarctic and Southern Ocean (ASO) species. **A.** Shared and unique species assessed by different extinction risk sources. **B.** Sources of assessments contributing to different taxonomic groups.

All sources of assessments focussed heavily on vertebrate species (Figure 2b). Nevertheless, including different sources and levels of assessments not only increased the number of species assessed but also the diversity of taxa and habitats assessed. NatureServe was a particularly important source of information for fungi assessments (98%), and through incorporating population-level assessments, the total number of invertebrate assessments increased to 676, compared with 428 assessments at the species level alone. Assessments published by the IUCN Red List were primarily for marine species at the species-level (59%) and terrestrial species at the population-level (42%) (Appendix S7). Assessments from national red lists were also primarily marine (68%). Conversely, NatureServe assessments were primarily for terrestrial species (52% of species- and 48% of population-level assessments).

Between the IUCN Red List and NatureServe there was a moderate but significant degree of agreement in standardised conservation statuses at the species level (K = 0.44, *n* = 315 species, *p* <0.01, z = 9.61, CI = 0.35 – 0.53). At the population-level, however, there was low, yet significant, agreement between the IUCN Red List and NatureServe (K = 0.11, *n* = 340 species, *p* <0.01, z = 4 .48, CI = 0.06 – 0.16). Confusion matrices showed the greatest differences between species listed as Threatened and Not Threatened for both species- and population-level assessments (Appendix S8). Furthermore, NatureServe provided species-level assessments for 12 species that were classified as Data Deficient by the IUCN Red List. Conversely, at the population level, the IUCN contributed information for 68 species listed as Data Deficient by NatureServe. Overall, integrating sources reduced the number of species with uncertain conservation status.

### Assessment Coverage

The first extinction risk assessments for ASO species were published in 1989 in a national red list by the Committee on the Status of Endangered Wildlife in Canada (Appendix S9). To date, there are a total of 5,403 assessments for 2,806 different species, which equates to 14% of known species in the ASO Species List with some form of assessment. Of the assessments with available dates (*n* = 1,725), 54% were published longer than a decade ago.

The proportion of ASO species assessed within taxonomic groups varied significantly (Figure 3). Vertebrates had the greatest proportion of species assessed (56%) despite them accounting for only 9% of total ASO species richness in the ASO Species List. Birds have the highest rates of assessed species overall (92%). Conversely, invertebrates accounted for 64% of ASO species richness, yet only 4% of invertebrate species have been assessed.

**Figure 3:**
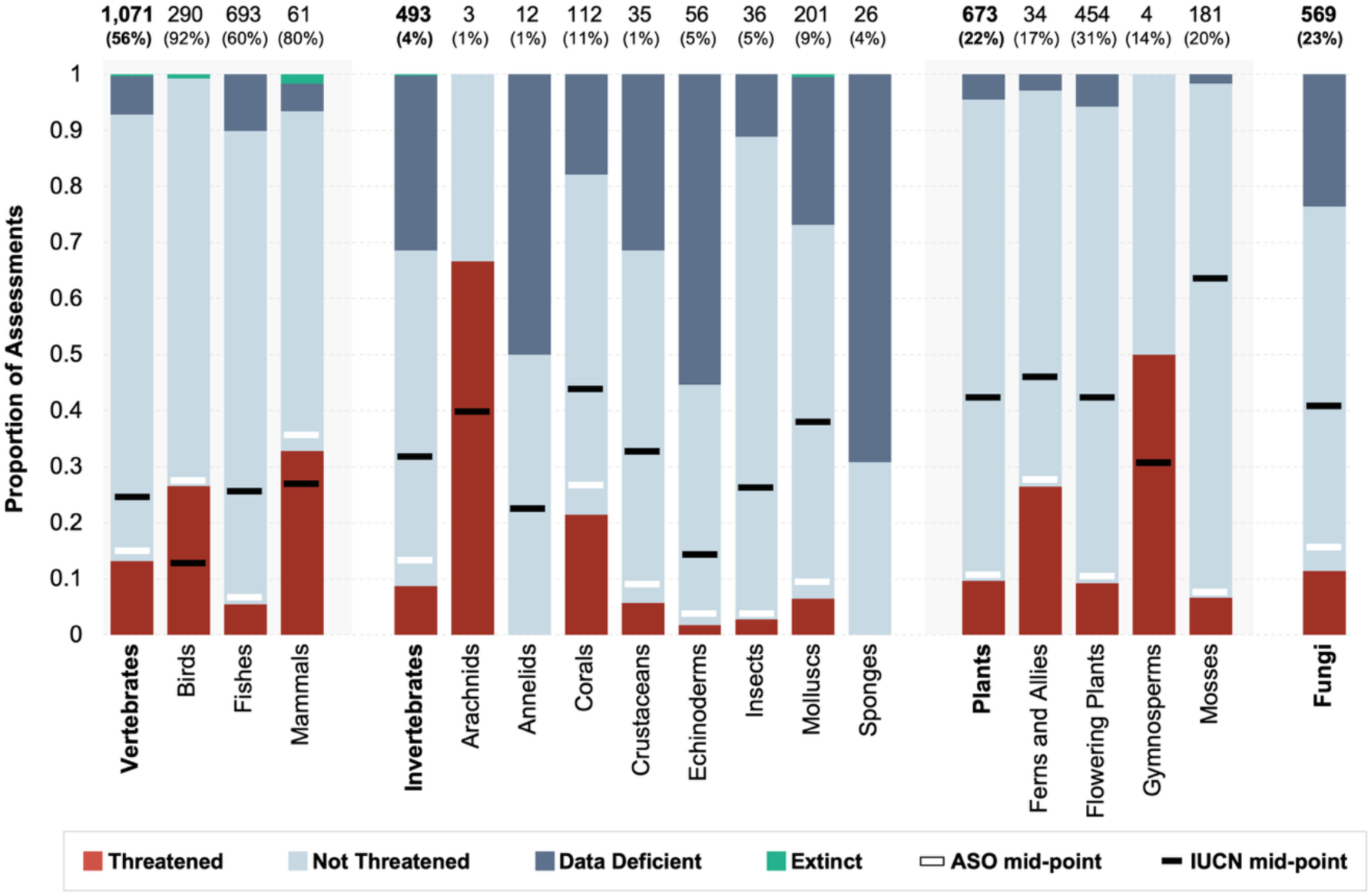
Threatened species in the Antarctic and Southern Ocean (ASO) by taxonomic group. The proportion of species in taxonomic groups (bold) and subgroups in each of the four standardised extinction risk categories. The total number of assessed species within each taxon is shown above the bar, and percentages in brackets show the proportion of known ASO species that have been assessed for each taxon. The white horizontal bars indicate the mid-point calculated as the number of extant species that would be considered Threatened if Data Deficient species are Threatened in the same proportion as Data Sufficient species (ASO mid-point). The black horizontal bars indicate the global taxon mid-point calculated using the IUCN summary statistics (IUCN, 2025).

Across all assessments, the proportion of species assessed from each of the three habitats did not match the proportion of ASO species occurring in those habitats (Appendix S10). Only 52% of marine species have been assessed, yet marine species contribute 73% of ASO species richness. Conversely, terrestrial and inland-water species account for a higher proportion of assessed species (40% and 8%, respectively) compared to their prevalence in the ASO (25% and 2%, respectively).

The geographic distribution of assessment density was strongly correlated with species richness (*rs* = 0.92, *n =* 7,177*, p* <0.01), indicating a positive monotonic relationship between the two per cell. In other words, regions with higher species richness also tend to have a greater number of species assessed for extinction risk. Assessment density was highly spatially variable (Figure 4a). Cells with high assessment density were concentrated around sub-Antarctic or Southern Ocean Islands, most notably Kerguelen Island (*n =* 566), South Georgia (*n =* 346), and Macquarie Island (*n =* 312). In contrast, the majority of terrestrial Continental Antarctica, and the Ross and Ronne ice-shelves, had cells with no assessments.

**Figure 4:**
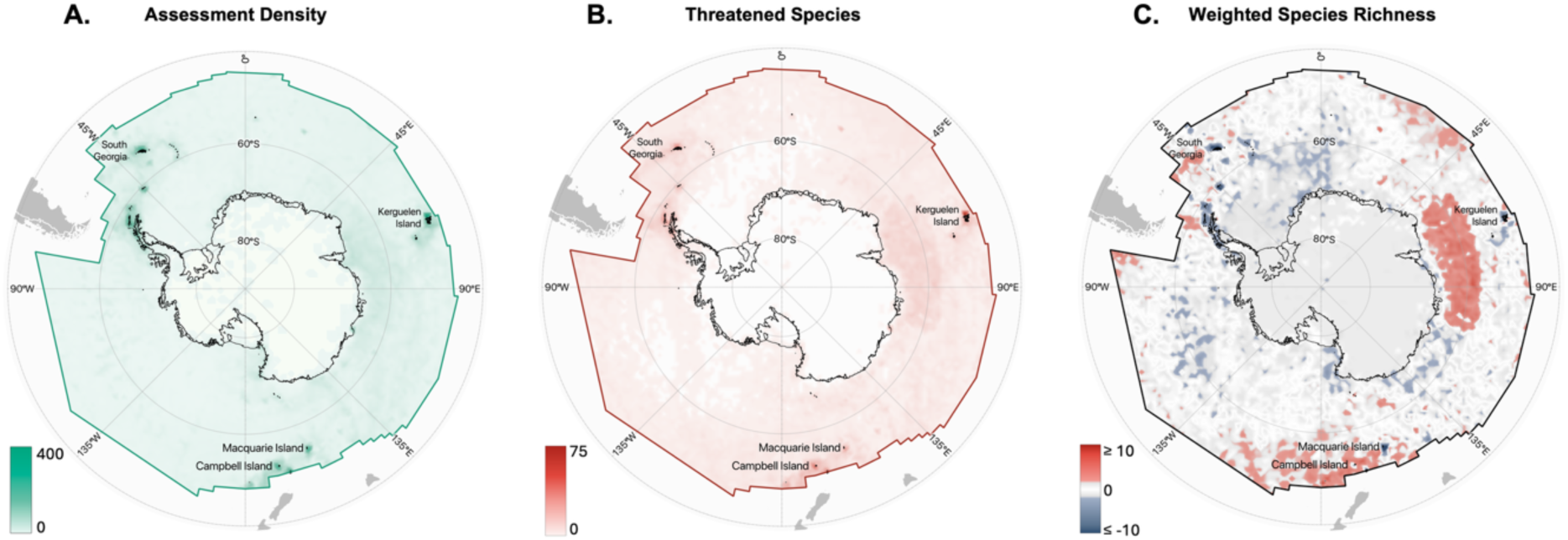
Geographic coverage of assessments and threatened species across the Antarctic and Southern Ocean. Cells are at a 100 km resolution**. A.** Assessment density measured as the total number of assessed species per cell. **B.** Threatened species richness. **C.** Weighted threatened species richness, i.e. the residuals of the relationship between the total number of Threatened species per cell and the total number of Data Sufficient species. Cells in red indicate positive residuals >2, with more Threatened species than expected when accounting for species richness and sampling effort. Grey cells represent residuals between +2 and -2, and cells in blue indicate negative residuals below <-2.

When examining assessment effort (the number of assessed species proportional to species richness per cell), many regions with high assessment numbers had only low to moderate assessment effort (Appendix S11). This indicates that, despite higher numbers of assessments, a substantial portion of species in these regions remained Not Assessed. For example, assessment effort revealed in cells surrounding Kerguelen Island, 19% of species are assessed, 15% around South Georgia, and 21% around Macquarie Island. Notably, 112 of 114 cells classified as having low assessment effort (between 0-10% of species assessed) were located within the Antarctic Treaty Region (south of 60°S). In contrast, cells with high assessment effort were generally those located in the Southern Ocean, away from landmasses. Notable Antarctic endemic species categorised as Not Assessed or Data Deficient include the insect *Belgica antarctica* (Jacobs, 1900), fungus *Cryomyces antarcticus* (Selbmann, de Hoog, Mazzaglia, Friedmann & Onofri, 2005), as well as all Collembola (springtail) and Acari (mite) species.

### Threatened species in the ASO

The proportions of Threatened species (Figure 3) and standardised residuals of assigned conservation status (Table 2) varied significantly across taxonomic groups. Note that the mid-point for arachnids and gymnosperms was not calculated because of low numbers of species assessed within the ASO (*n* 7 10). Chi-squared tests revealed a significant association between conservation status assignment and both major taxonomic groups (χ² = 268.74, df = 6, *p* <0.01) and taxonomic sub-groups (χ² = 525.98, df = 32, *p* <0.01), indicating some taxa are disproportionately represented in certain conservation statuses.

**Table 2:**
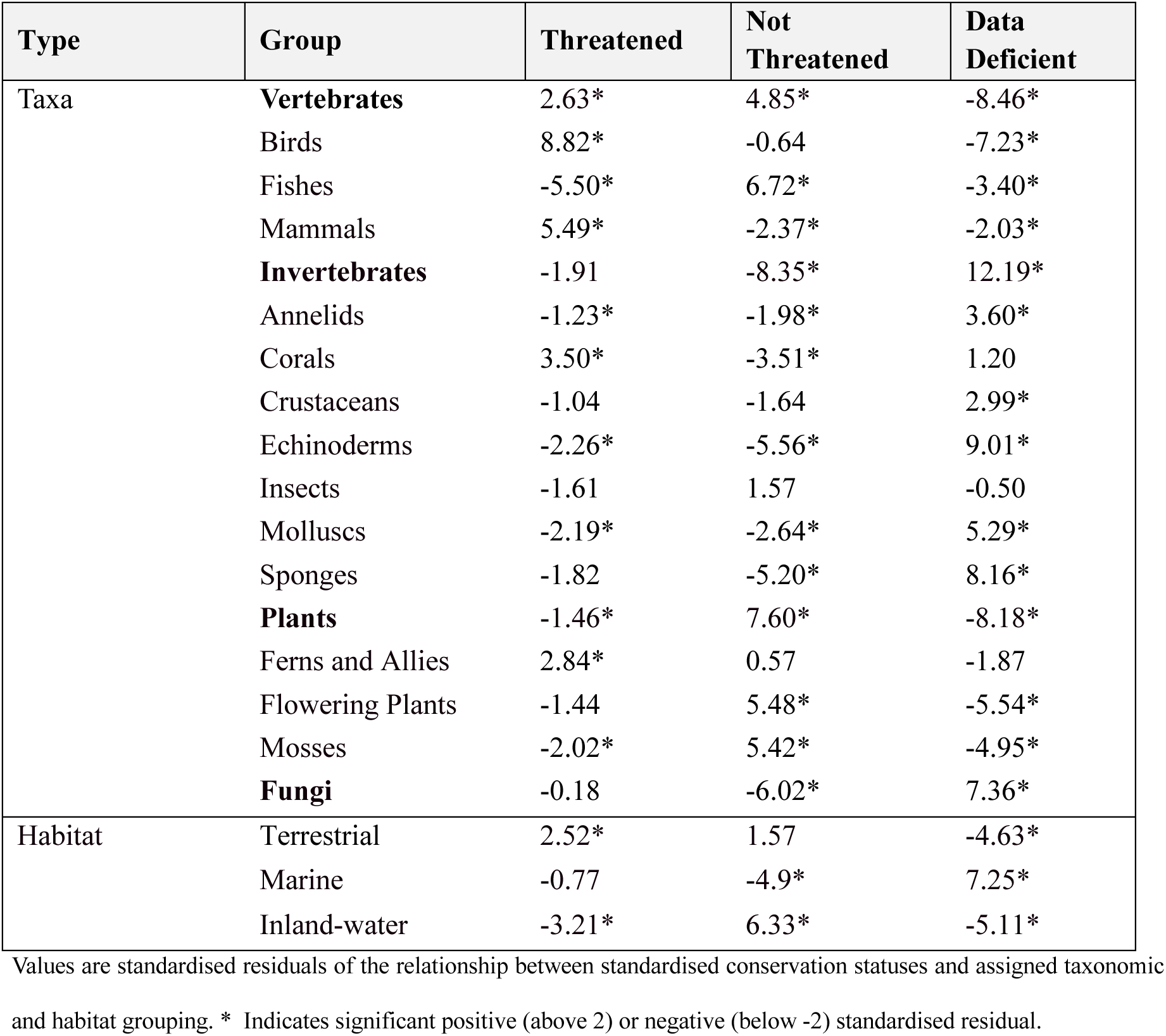
The prevalence of taxonomic and habitat groups in standardised conservation statuses (Threatened, Not Threatened, Data Deficient) for species assessed for extinction risk within the Antarctic and Southern Ocean.

**Table 3:** The proportions of Antarctic and Southern Ocean species impacted by different drivers of population decline for different standardised conservation statuses (Threatened, Not Threatened, Data Deficient), in comparison with global proportions derived from the IUCN Red List (IUCN, 2025).

| Driver | Threatened | Not Threatened | Data Deficient | Global |
| --- | --- | --- | --- | --- |
| 1: Residential and commercial development | 0.26 ↓ | 0.22 ↓ | 0.04 ↓ | 0.29 |
| 2: Agriculture and aquaculture | 0.24 ↓ | 0.14 ↓ | 0.08 ↓ | 0.56 |
| 3: Energy production and mining | 0.10 ↓ | 0.07 ↓ | 0.02 ↓ | 0.16 |
| 4: Transportation and service corridors | 0.19 ↑ | 0.14 ↑ | 0.02 ↓ | 0.10 |
| 5: Biological resource use | 0.77 ↑ | 0.70 ↑ | 0.87 ↑ | 0.48 |
| 6: Human intrusions and disturbance | 0.23 ↑ | 0.20 ↑ | 0.00 ↓ | 0.07 |
| 7: Natural system modifications | 0.13 ↓ | 0.09 ↓ | 0.04 ↓ | 0.27 |
| 8: Invasive and problematic species, genes, and diseases | 0.57 ↑ | 0.33 ↑ | 0.08 ↓ | 0.18 |
| 9: Pollution | 0.33 ↑ | 0.27 ↑ | 0.08 ↓ | 0.16 |
| 10: Geological events | 0.01 | 0.01 | 0.00 ↓ | 0.01 |
| 11: Climate change | 0.49 ↑ | 0.36 ↑ | 0.09 ↓ | 0.17 |
| 12: Other | <0.01 | <0.01 | 0.00 ↓ | <0.01 |
| Total number of species | 195 | 513 | 53 | 97,116 |

Data are shown using the conservation statuses from the IUCN Threat Classification Scheme (version 3.3) for 775. Note that a species may be listed under more than one threat. Arrows indicate when the result is higher or lower than the global proportion calculated by the IUCN.

Among taxonomic subgroups, mammals had the highest proportion of Threatened species, with over one-third classified as Threatened (mid-point: 35%, range: 33-38%), followed by birds (mid-point: 27%, range: 27-27%). Standardised residuals supported these patterns, with significantly more Threatened mammal and bird species than expected. Threatened species included the sub-Antarctic endemic Tristan Albatross (*Diomedea dabbenena* Mathews, 1929) and the New Zealand Sea Lion (*Phocarctos hookeri* Gray, 1844). Notably, mammals and birds were the only two taxa with higher proportions of threatened species in the ASO compared to global estimates (Figure 3), as published by the IUCN Red List (IUCN mammal mid-point: 27%, IUCN mammal range: 23-36%; IUCN bird mid-point: 12%, IUCN bird range: 12-12%) (IUCN, 2025).

Invertebrates generally had low proportions of Threatened species, but a high amount of uncertainty, shown by the large range (mid-point: 13%, range: 9-40%). Among invertebrate sub-groups, corals were the most Threatened (mid-point: 26%, range: 21-39%) (Figure 3), consistent with their significant positive standardised residual (Table 2). Threatened corals include the deep-sea species *Umbellula lindahli* Kölliker, 1875. Overall, the proportion of Threatened invertebrates within the ASO was substantially lower than global averages (IUCN invertebrate mid-point: 31%) (IUCN, 2025).

Among plants and Fungi, 10% of ASO plant species (range: 10-14%) along with 15% of Fungi (range: 11-35%) were classified as Threatened. Standardised residuals for plants and Fungi suggest no significant deviation from expected proportions. However, these mid-point values are considerably lower compared to global IUCN estimates for plants (mid-point: 42%) and Fungi (mid-point: 40%) (IUCN, 2025).

More than one-third of ASO invertebrate species were classified as Data Deficient. Invertebrates showed particularly high levels of uncertainty in the echinoderms (55% Data Deficient) and sponges (69% Data Deficient), supported by the high positive standardised residual for Data Deficient invertebrates (Table 2). Likewise, Fungi showed substantial uncertainty, with 24% of species classified as Data Deficient and a significant positive standardised residual. In contrast, vertebrates and plants had only 7% and 4% of Data Deficient species, respectively, and significantly low negative standardised residuals. This demonstrates their underrepresentation in the Data Deficient category.

Chi-squared tests revealed a significant association between conservation status and habitat type (χ² = 78.66, df = 4, *p* <0.01). Terrestrial species were overrepresented in the Threatened category, with a significant negative standardised residual (Table 2) and 46% of species classified as Threatened (Appendix S10). Terrestrial species were underrepresented in the Data Deficient category, showing significant negative standardised residuals. In contrast, marine species showed the expected proportion of Threatened assessments but were overrepresented in the Data Deficient category, with significant positive standardised residuals.

The highest concentrations of Threatened species per cell were recorded in the sub-Antarctic and maritime Antarctic regions (Figure 4d), particularly on Kerguelen Island (*n* = 75), Campbell/Motu Ihupuku Island (*n* = 43), and South Georgia (*n* = 41). However, the weighted Threatened species richness (Figure 4e) for these cells had significantly low standard residuals (between -3.1 and -12.4). This indicates that despite high absolute numbers of Threatened species, these areas contain fewer Threatened species than expected when accounting for overall species richness and assessment density.

### Threats to Species

For ASO species, ‘Biological Resource Use’ was the most frequent threat affecting 77%, 70%, and 87% of Threatened, Not Threatened, and Data Deficient species, respectively (Table 4). The IUCN (2025) defines ‘Biological Resource Use’ as a threat from “consumptive use of ‘wild’ biological resources, including both deliberate and unintentional harvesting effects”. This encompasses activities such as hunting, collecting, and/or fishing. Globally, ‘Biological Resource Use’ is a major threat affecting 48% of assessed species globally, the second-highest proportion among all conservation statuses (IUCN, 2025). For Threatened species, ‘Invasive and problematic species, genes, and diseases’ was listed as the second major threat, affecting 57% of ASO Threatened species, followed by ‘Climate Change’ (49%). ‘Invasive and problematic species, genes, and diseases’ includes threats which may arise from alien species, introduced genetic material, and diseases (IUCN, 2025). Both ‘Invasive and problematic species, genes, and diseases’ and ‘Climate Change’ are overrepresented in ASO species in comparison with global populations, where they only affect 18% and 17% of IUCN assessed species, respectively (IUCN, 2025).

**Table 4:** Key management actions to strengthen the information needed to manage and measure species conservation progress in the Antarctic and Southern Ocean, including to meet Target 4 and Goal A of the Kunming Montreal – Global Biodiversity Framework (GBF) and to effectively implement the Antarctic Specially Protected Species framework.

| Management Action | Description |
| --- | --- |
| 1. Continued development of comprehensive species checklists and occurrence data. | Up-to-date inventories on species and their distribution are foundational for completion of extinction risk assessments. This requires coordinated efforts to consolidate data from diverse sources, systematic incorporation of new research and remote sensing approaches, and adoption of standardised data sharing frameworks. |
| 2. Integration of existing extinction risk assessments from multiple sources and multiple scales. | Integration maximises the current utility of extinction risk assessments when information availability is low and benefits understanding species intraspecific extinction risk variability. This can be achieved through developing consistent methodologies and imputation techniques to improve comparability amongst sources and scales. |
| 3. Strategic prioritisation of future assessments and re-assessments. | This should account for current taxonomic, geographic, and temporal biases, moving beyond the current ‘ad hoc’ conservation approach which has historically targeted charismatic, yet not necessarily ecologically significant, species. |
| 4. Alignment of conservation management with Threatened species protection. | Explicit consideration of Threatened species in protected area designation and increased use of the Antarctic specially protected species would establish actions towards achieving GBF and helping Antarctic conservation management transition from a reactive to a more proactive position. |

### Protected Area Coverage of Threatened Species

There was no consistent relationship between assigned conservation status and the proportion of species extent of occurrence overlapping with marine (χ^2^ = 0.49, df = 3, *p* = 0.92, ε^2^ = 0) species and protected areas (Appendix S12 and S13). For terrestrial and inland-water species (χ^2^ = 16.63, df = 3, *p* <0.01, ε^2^ <0.01), differences in protected area overlap exist between species that were assessed compared to Not Assessed, but no difference was seen between the standardised conservation statuses. Species classified as Extinct were not included in this analysis because of low numbers. The ASO species with no protected area coverage included the Threatened Chatham Island Petrel, *Pterodroma axillaris* (Salvin, 1893).

## Discussion

Region-specific overviews of extinction risk knowledge are essential for understanding how biodiversity is responding to threats and environmental change (Kuipers et al., 2019). This first regional synthesis of extinction risk for Antarctic and Southern Ocean (ASO) species reveals significant taxonomic, habitat, and geographic bias, along with temporal shortfalls in assessment coverage, limiting the capacity for conservation management to identify and monitor species vulnerability with sufficient warning time for action (Stanton et al., 2015). Notably, despite mammals and birds being proportionally more threatened in the ASO than globally, protected areas in the region do not provide greater coverage for Threatened species compared to Not Threatened or Not Assessed species. By integrating assessments across sources and scales (ranging from population- to species-level assessments), our data-inclusive approach reduced knowledge gaps compared with using any single source or scale, demonstrating that no individual assessment database or framework provides a complete picture of the region. Nevertheless, the limited representativeness of available assessments still significantly constrains the evaluation of progress towards the Kunming-Montreal Global Biodiversity Framework (GBF) in the ASO and complicates the designation of Antarctic Specially Protected Species (SPS) based solely on International Union for the Conservation of Nature (IUCN) Red List extinction risk.

### Taxonomic and Geographic Assessment Bias

Bias towards assessments of vertebrates aligns with well-documented regional (Jones et al., 2025) and global patterns in which conservation research and extinction risk assessments disproportionately favour animals and charismatic megafauna (Alfonzetti et al., 2020), while neglecting invertebrates (Cardoso et al., 2011; Martín et al., 2010). This bias is particularly problematic in the ASO where invertebrates account for much of community species richness (Clark et al., 2015). As a result, conservation decisions generally rely on a narrow set of well-known species as proxies for broader biodiversity (Clark & May, 2002), despite growing evidence that using well-studied and charismatic species as ‘umbrella’ or ‘surrogate’ proxies often provides poor representation for overall biodiversity (Cardoso et al., 2011; Tälle et al., 2023). This is especially pronounced in systems dominated by narrow-ranged species (Yang et al., 2023), such as many of the terrestrial ASO species, where adequate assessments for certain taxa exist alongside neglected groups.

Notably, the IUCN have reported that 100% of global bird species have an extinction risk assessment, based on species lists from Birds of the World and BirdLife International (IUCN, 2025). However, we identified 24 bird species present within the ASO (8% of the total number of ASO species) that lack any form of extinction risk assessment. These include the sub-Antarctic endemic Crozet Shag (*Phalacrocorax melanogenis* Blyth, 1860), Eastern Rockhopper penguin (*Eudyptes filholi* Hutton, 1879), and the Gibsons Albatross (*Diomedea gibsoni* Robertson & Warham, 1992). While it is important to acknowledge the considerable effort in assessing 92% of ASO birds, these gaps further highlight the need for targeted attention to the ASO avifauna, given the ecological endemism and uniqueness of the region’s bird communities (Dehling & Chown, 2025).

The strong correlation between species and assessment richness may be a product of activity and research being completed in well-established areas, such as research stations, thereby reinforcing geographic biases in both species data and assessment coverage (Patterson et al., 2025). However, low assessment effort present in locations such as the sub-Antarctic indicates that biodiversity hotspots, such as the sub-Antarctic islands remain underassessed, suggesting that these regions may be overlooked in conservation management and/or under-resourced for biodiversity research.

### Outdated Assessments

A high proportion of ASO assessments were over 10 years old, reducing likely accuracy and usefulness (Rondinini et al., 2014). The reliability of assessment data decreases over time (Rondinini et al., 2014), and due to the changing nature of taxonomy, older assessments may additionally be unrepresentative of current taxonomic classification (Clarke et al., 2025; Lončarević et al., 2024). As a result, IUCN assessments are officially classed as “outdated” after 10 years since publication (IUCN, 2025). The proportion of assessments in the ASO exceeding this threshold was more than three times higher than the reported global average of outdated IUCN assessments (17%) (Rondinini et al., 2014). Basing management decisions on outdated assessments could lead to misallocation of resources (Rondinini et al., 2014) and may obscure early warning-signals of ecosystem change. The high proportion of outdated assessments in the ASO likely reflects the more opportunistic, as opposed to strategic, approach to assessment completion, compounded by the absence of a coordinated framework for reassessments. Up-to-date extinction risk assessments are essential for providing accurate warning times for detecting species declines and for monitoring trends in species status (Glasnović et al., 2024; Rodrigues et al., 2006). This is amplified in ASO where the rates of environmental change are among the fastest globally, increasing the rate at which assessment data becomes outdated and unreliable (Gorodetskaya et al., 2023; Morley et al., 2020; Nel et al., 2023).

### Threatened Species

Mammals and birds were the only taxa with a higher proportion of Threatened species in the ASO than globally, as reported by the IUCN Red List, with 27% for mammals and 12% for birds (IUCN, 2025). This contrasts with the long-standing paradigm that birds are more Threatened in tropical latitudes (Lees et al., 2022; Reif & Štěpánková, 2016) and provides evidence to support predictions of declining Antarctic seabird trajectories (Lee et al., 2022). Elevated threat levels of ASO birds and mammals may be a reflection of species range-restriction, trait distinctiveness (Dehling & Chown, 2025), climate-driven shifts in prey availability (Bestley et al., 2020), and invasive species impacts in important breeding grounds (Dilley et al., 2018; Tasker et al., 2000). This higher proportion of Threatened mammal and bird species in the ASO, despite these taxa dominating assessments, suggests a misalignment between extinction risk assignment and the allocation of conservation resources (Guénard et al., 2025) and an underutilisation of the Antarctic Specially Protected Species (SPS) designations (Hughes et al., 2024) (Table 4).

Despite high numbers of Threatened species on the sub-Antarctic islands and the Antarctic Peninsula, these locations had fewer Threatened species than expected after accounting for assessment density and species richness. This pattern may indicate either that these species-rich areas are less exposed to threatening processes, or that species on these islands have lower inherent vulnerability to extinction compared with global patterns (Kier et al., 2009). Features of these island ecosystems such as high insularity, has been reported to be a strong determinant of high extinction risk, with island species typically more vulnerable due to high endemism, range restriction, and narrower niche breadth (Fernández-Palacios et al., 2021; Gaston, 2009; Howard et al., 2020; Kier et al., 2009). Additionally, current uneven extinction risk assessment coverage and high may mask true extinction risk in these regions. Nevertheless, these results suggest that sub-Antarctic islands and the Antarctic Peninsula may function as refugia for some taxa, emphasising the importance of targeted conservation measures to safeguard endemic and Threatened species. As, additional data and assessments becomes available, the proportion of Threatened species is expected to change, particularly for poorly assessed taxa such as invertebrates, where many Data Deficient species may likely prove Threatened (Borgelt et al., 2022).

The current ASO protected area network provides no greater coverage for Threatened species than for Not Threatened or Not Assessed species in either marine or terrestrial realms, with negligible effect sizes. The lack of protected area coverage for the Threatened Chatham Island Petrel exemplifies this incongruence between conservation status and area-based protection. This may, in part, reflect that not all protected areas are designated primarily for biodiversity conservation, but rather for their scientific, historic, aesthetic, or wilderness values (Antarctic Treaty Secretariat, 2025). This is consistent with growing criticism of the current protected area system for limited ecological representativeness, insufficient area coverage, and poor management effectiveness (Brooks et al., 2020; Coetzee et al., 2017; Hughes et al., 2016). Our findings support these concerns and underscore the need for the establishment of protected areas that explicitly prioritise Threatened biodiversity, in conjunction with well-designed management plans and conservation measures (Kearney et al., 2020) (Table 4) which account for future changes in species range dynamics and habitat suitability (Hughes et al., 2021) (Table 4). As assessment coverage increases, particularly for invertebrates and sub-Antarctic species, the true extent of this incongruence is likely to increase.

### Integration of species- and population-level assessments

Incorporating population-level data alongside species-level assessments is critical for two reasons. First, extinction risk assessments are resource-intensive, and the lack of population and abundance data often impedes their completion (Kozlowski, 2008; Zamin et al., 2010). By integrating population-level assessments, we were able to leverage additional data in regions where information is relatively scarce. This approach added significant value by capturing taxa and habitats that would otherwise be overlooked, thereby providing a more comprehensive representation across ASO biodiversity. The second reason is that threats are rarely uniformly distributed across species’ ranges. Population-level assessments can reveal localised trends and specific pressures affecting population trajectories, providing critical warning on species population change that may precede species-level reclassifications (Kuipers et al., 2019). Furthermore, conservation action typically occurs at the population level accounting for intraspecific variation in extinction risk assessments is essential for evaluating effectiveness of implemented management strategies (Cowlishaw et al., 2009). Consistent with previous studies (Cowlishaw et al., 2009; Holz et al., 2022; Mounce et al., 2018; Zamin et al., 2010), our findings support that there is much to be gained by integrating extinction risk information across multiple spatial levels and sources to strengthen conservation planning.

A major challenge in applying extinction risk results and integrating different assessment sources is the presence of conflicting conservation statuses. Variability in status between frameworks is expected and arises from differences in abundance estimates between assessed populations, assessment objectives, bureaucratic constraints, protocol misapplication, or disparities in financial or technical resources (Brito et al., 2010; Possingham et al., 2002; Zamin et al., 2010). Additionally, when comparing extinction risks at different scales, species-scale assessments may not always align with population-scale trends (Glasnović et al., 2024). Regardless of the underlying cause, inconsistencies in conservation status for the same species undermine the credibility of extinction risk assessments (Brito et al., 2010), and complicate their application in policy frameworks, including tracking progress against GBF. These challenges may be particularly pronounced in the ASO, where obtaining timely and detailed information is difficult and species distributions often span multiple international and geopolitical jurisdictions. Consequently, species that occur entirely within global commons or the Antarctic Treaty System may be overlooked, or they may hold assessments under multiple frameworks that are not directly comparable (de Villiers et al., 2005). To address these challenges, the development and adoption of standardised assessment protocols within the ASO or the consistent application of assessment integration, would be highly beneficial (Table 4). While our results depend on the process used to standardise and prioritise species, they demonstrate the potential of this approach and highlight the need for further research to refine and expand standardised assessment methods in the future.

## Conclusion

This analysis represents an important step in establishing a baseline of Antarctic and Southern Ocean (ASO) extinction risk to enable monitoring of biodiversity trends under ongoing environmental change. Yet systemic gaps and biases in extinction risk assessments, and a mis-match between Threatened species and protected area coverage, significantly constrains current conservation management. At a time when global species declines demand considered and timely action, a strategically prioritised approach to extinction risk assessments and better protection for species most at risk will not only improve regional conservation outcomes but also contribute to a more accurate global understanding of progress towards international biodiversity targets.

## Acknowledgements

We would like to thank Dr. Samuel G. Beale for his useful comments on the draft manuscript, Caitlin A. Selfe for her help with QGIS, and Dr. David A. Clarke for his coding assistance. This research is supported by ARC SRIEAS Grant SR200100005 Securing Antarctica’s Environmental Future (SAEF) and by the Commonwealth through an Australian Government Research Training Program Scholarship (DOI: https://doi.org/10.82133/C42F-K220).

