## Supplementary material for "Revealing the Widespread Bias of Extinction Risk in the Antarctic and the Southern Ocean"

Madison G. Farrant[1], W. P. Amy Liu[1], Melodie A. McGeoch[1]
[1]Securing Antarctica's Environmental Future, School of Biological Sciences, Monash University, Melbourne 3800, Victoria, Australia

This PDF file includes:

**Appendix S1:** Detailed methods for data acquisition, processing, and analysis.

**Appendix S2:** National and supra-national publications of population-level extinction risk assessments for species within the Antarctic Southern Ocean downloaded through the National Red List Database.

**Appendix S3:** Lexicon standardisation of different sources of extinction risk assessments to five conservation statuses (Extinct, Threatened, Not Threatened, Not Assessed, Data Deficient) where the standardised extinction risk category differed from the original category.

**Appendix S4:** Workflow used to prioritise Antarctic and Southern Ocean species extinction risk conservation statuses where multiple assessments for a species were present.

**Appendix S5:** Fields present within the Occurrence Dataset, Antarctic and Southern Ocean (ASO) Species List, and the Extinction Risk Datasets.

**Appendix S6:** Current protected areas found either partly or wholly within the Antarctic and Southern Ocean Marine Ecosystem Assessment of the Southern Ocean (MEASO) region.

**Appendix S7:** Habitat coverage of extinction risk assessments by source for assessments conducted at the species- and population-levels.

**Appendix S8:** Confusion matrix results of extinction risk agreement between different sources and scales of extinction risk assessments.

**Appendix S9:** Cumulative publication dates of extinction risk assessments for Antarctic and Southern Ocean Species (ASO).

**Appendix S10:** Distribution of extinction risk assessments and conservation statuses for Antarctic and Southern Ocean (ASO) species across different habitats (marine, terrestrial and/or inland-water).

**Appendix S11:** Geographic coverage of assessment effort for Antarctic and Southern Ocean Species.

**Appendix S12:** The overlap between the extent of occurrence of species distributions (EOO) and terrestrial (including Inland-water habitats) and marine Antarctic and Southern Ocean protected areas for each standardised conservation statuses.

**Appendix S13:** Dunn's test *p*-value results between the overlap between the extent of occurrence of species distributions and terrestrial and marine Antarctic and Southern Ocean protected areas for each standardised threat category.

**References**

**Appendix S1:** Detailed methods for data acquisition, processing, and analysis.

**Dataset Acquisition and Development**

The following methods were used for the compilation of three datasets, the 'Occurrence Dataset', the 'ASO (Antarctic and Southern Ocean) Species List', and the 'Extinction Risk Dataset'. All data acquisition and analysis were conducted using RStudio version 2024.04.0+735 (R Core Team, 2022) with code used available via FigShare (https://doi.org/10.6084/m9.figshare.31175305) (Farrant et al., 2026). The region defined as the ASO in this research follows that adopted in the Marine Ecosystem Assessment for the Southern Ocean (MEASO) (Constable et al., 2023).

*Occurrence Dataset*

The lack of knowledge of described species is a significant challenge in Antarctica (Vecchi et al., 2025). This current knowledge shortfall is largely attributed to the challenging conditions, relatively short history of Antarctic science, and presence of many cryptic species, which leaves many aspects of Antarctic and the surrounding Southern Ocean relatively data poor (Pertierra et al., 2025; Pertierra et al., 2023). As a result, there is no comprehensive species checklist for the MEASO region. Therefore, we have used occurrences as the foundation for identifying species present within the ASO, and for geospatial analysis of extinction risk trends.

Extant species occurrence records were downloaded from four open-access databases for species within the Antarctic and Southern Ocean (ASO). Databases used were: the Global Biodiversity Information Facility (GBIF) (Gbif.org, 2025a, 2025b, 2025c), accessed using the *rgbif* package with the 'occ_download()' function (Chamberlain, 2017); Ocean Biodiversity Information System (OBIS) (), accessed using the *robis* package and the 'occurrence()' function (Provoost et al., 2022); the Biodiversity of ice-free Antarctica database version 5 (Terauds et al., 2025b), downloaded through the Australian Antarctic Data Centre (Terauds et al., 2025a); and the SIVFLORA database of sub-Antarctic vascular flora version 1.0 (Guerrero et al., 2025), accessed via Zenodo (Guerrero & Medina, 2025). All occurrence data were downloaded between 21st of January 2025 and 11th of March 2025. Presence only occurrence records were retained if they met the following criteria: (1) georeferenced within the MEASO region with no reported geospatial issue; (2) classified within the kingdoms Animalia, Fungi, or Plantae; (3) identified to the species level (i.e. the 'taxonRank' data field was "species"); and (4) listed in the 'basisOfRecord' with "observation", "human observation", "machine observation", "material sample", "occurrence", or "material citation". The initial downloads comprised 8,429,630 occurrence records, including both taxonomically accepted and synonym species.

Scientific names and higher taxonomic classifications (kingdom, phylum, class, order, family, and genus) were computationally harmonised using GBIF Backbone Taxonomy (GBIF Secretariat, 2023) through the name_backbone() function of the *rgbif* package (Chamberlain, 2017). Taxonomic harmonisation ensures standardisation and reproducibility of taxonomic names, in addition to resolving any synonyms. For the harmonisation process scientific names were pre-processed to remove name authorship. Fuzzy matching was not used. Original species names recorded verbatim from source databases were preserved in the 'originalNameUsage' field to facilitate cross-referencing with historical sources. Records with misspelled or unrecognised species names that could not be harmonised computationally, were manually corrected using GBIF Backbone Taxonomy (GBIF Secretariat, 2023) or excluded from the dataset ($n$ = 115 records) (or example, "*Octopoxenus antarcticus*"). A further 3,094 records were removed that had insufficient identification at species level (e.g. listed with a 'taxonRank' as 'species' however, identified as "*Grimmia sp.*"). Duplicate records, defined as those with matching information across the 'scientificName', 'basisOfRecord', 'decimalLatitude', 'decimalLongitude', and 'eventDate' columns were removed ($n$ = 1,595,713 records). The *coordinatecleaner* package was used to detect and exclude any potentially erroneous coordinates. Coordinates that corresponded to capitals, centroids, zeros, or statistical outliers using the median absolute deviation method using the clean_coordinates() function were flagged (Zizka et al., 2023). The 11,701 flagged records were subsequently removed. Occurrences located south of 60° latitude fall within the Antarctic Treaty System (ATS) management area and are therefore the responsibility of the Consultative parties to the Antarctic Treaty (Saul & Stephens, 2015). In contrast, areas north of 60° latitude, fall either within national jurisdiction (such as the sub-Antarctic Islands) or are considered part of the global commons (de Villiers et al., 2005). To distinguish between these regions, 'higherGeographyID' was used to classify records as either within the ATS (labelled "Antarctic") or outside it (labelled "sub-Antarctic). Where occurrences were extracted form GBIF the 'gbifID' column was retained for cross-referencing of information.

The final Occurrence Dataset (Occurrence_dataset.csv) contained 6,819,007 species-level occurrences from the ASO region. This dataset contains an inventory of marine, terrestrial, and inland-water dwelling species, with observation dates spanning 1697 to 2025. Each record includes information on species taxonomy, occurrence status, observation date, and data source. Additional fields related to species habitat and extinction risk were subsequently incorporated after the development of the Extinction Risk Dataset.

*Antarctic and Southern Ocean Species List*

To supplement the species within the Occurrence Dataset, additional taxa were incorporated using the TerrANTALife species checklist (Pertierra et al., 2024). TerrANTALife provides a list of Antarctic terrestrial and inland-water species, which spans all Antarctic living and viral organisms. The TerrANTALife eukaryotic checklist version 1.0 was downloaded through Digital CSIC on the 21st of January 2025 (Pertierra et al., 2023). All records were taxonomically harmonised as above using GBIF Backbone Taxonomy (GBIF Secretariat, 2023), and filtered for those which met the criteria as outlined for the occurrence downloads above. TerrANTALife species were cross checked with those already present within the Occurrence Dataset. Among the animal, plant, and fungal species listed within TerrANTALife, 78% had corresponding occurrence records in the Occurrence Dataset. The remaining 356 species were then integrated with Occurrence Dataset species to create the ASO Species List. Any species which was incorporated into the ASO Species List from TerrANTALife, had 'TerrANTALife' added to the 'datasetName' field. Additionally, each species in the ASO Species List also present within the TerrANTALife checklist had the corresponding 'taxonID' added for cross-referencing to the original dataset.

The final ASO Species List (ASO_species_list.csv) contained 20,764 species spanning 42 phyla. As with the Occurrence Dataset, the ASO Species List also compiled information on species taxonomy and source. Information regarding species habitat and extinction risk was subsequently added to the ASO Species List after the development of the Extinction Risk Dataset.

*Extinction Risk Dataset*

Extinction risk data were extracted from literature and four databases. Namely the IUCN Red List of Threatened Species (henceforth, the IUCN Red List) (IUCN, 2025a), NatureServe (NatureServe, 2025), the Convention on International Trade in Endangered Species (CITES) (CITES, 2025), and the National Red List Database (ZSL and IUCN National Red List Working Group, 2022). The National Red List Database compiles extinction risk assessments published by both national and governmental conservation authorities. All database downloads of extinction risk assessments were made between the 7th of February 2025 and the 19th of February 2025. To ensure comprehensiveness, additional extinction risk assessments and population viability analyses were augmented through literature searches in Scopus (Elsevier, Amsterdam, Netherlands) and in the Web of Science Core Collection (Clarivate Plc, Pennsylvania, United States) on the 28th of January 2025 and the 16th of March 2025, respectively. The Boolean search string ("*Southern Ocean*" OR *Antarctic** OR "*sub-Antarctic*") AND ("*population viability analysis*" OR "*extinction risk*" OR "*Conservation status**") was used to search publication title, abstract, and keyword fields. There were 176 unique papers published between 1981 and 2025 that were further screened for relevant information. One of which (Padin & Calviño, 2023) was unable to be sourced beyond the abstract. Information was deemed relevant if an independent assessment of extinction risk was completed for species on the ASO Species List. Thirty assessments from literature were retained for 28 unique species on the ASO Species List from seven different publications.

For all extinction risk assessments, species names were taxonomically harmonised using GBIF Backbone Taxonomy (GBIF Secretariat, 2023) as above and filtered for species within the ASO Species List. Information on the database or literature source was retained in the 'assessmentSource' field. For extinction risk assessments downloaded through the National Red List Database, assessments were derived from 61 separate publications with the publication title added to the 'assessmentPublication' field. When available, information was also retained from assessments regarding the year of assessment and assigned habitat. The scale of each assessment was also classified as either being at a species-level (global assessments), or a population-level (national to sub-global assessments) within the 'assessmentScope' field.

To support a comprehensive analysis of taxonomic and habitat coverage in extinction risk assessments and threatened species, all ASO species were assigned habitat and taxonomic groupings for comparison. Habitat information was obtained from multiple data sources throughout data acquisition. Priority was given to habitat data when provided by species assessment sources, such as the IUCN Red List, to enable comparisons with global assessments of habitat and extinction risk. For species lacking habitat information from their extinction risk source, habitat data from the Register of Antarctic Species (RAS, 2025) were matched to the corresponding taxonomically harmonised name. Remaining habitat gaps were addressed by assigning habitats based on species' higher taxonomic classification for taxonomic groups which are known to be entirely marine or terrestrial. Finally, remaining gaps were assigned using the Occurrence Dataset to identify geographic coordinates that intersected with either terrestrial or marine regions of the *rnaturalearth* medium-scale landmass polygons (Massicotte et al., 2023).

Major taxonomic groupings were assigned at the kingdom level, with Animalia further separated into 'vertebrates' (phylum Chordata) and 'invertebrates' (non-Chordata). Taxonomic sub-groups, such as 'fishes', 'mammals', and 'mosses', were assigned in accordance with the IUCN summary statistics for comparison (IUCN, 2025a). The additional taxonomic subgroups of 'sponges', 'echinoderms', and 'annelids' were added based on species phylum-level taxonomy (i.e. phyla Porifera, Echinodermata, and Annelida, respectively). It should be noted that the IUCN taxonomic categories classify fungi and protists under a single taxonomic group, however, for the purpose of this study as no protists were included, we refer this category as 'Fungi'. Both taxonomic groupings in the 'taxonCategory' and 'habitat' fields were added to the corresponding species within the Occurrence Dataset and ASO Species List.

To address variability in the lexicon used between assessment sources, assessment statuses were standardised into five conservation statuses: Extinct, Threatened, Not Threatened, Not Assessed, and Data Deficient. The original criteria used by each source was reviewed to ensure that the standardised threat category accurately reflected the intended conservation status. Assessments from the

Recopilación de la Información Sobre la Biodiversidad de Honduras were excluded from further analysis as the terminology used could not be confidently reconciled. Standardised assessments were added to the 'standardisedAssessment' field, with the original assessment categories as assigned by the source, retained in the 'verbatimAssessment' field. For subsequent data analyses on Threatened species, in the instances where species had multiple conservation statuses, priority was given to the threat category of assessments completed within the last 10 years, then was conservatively assigned to the most threatened category. Most threatened categories ranked from highest to lowest were: Extinct, Threatened, Not Threatened, Data Deficient, and Not Assessed. The prioritised assessments were added to the 'selectedAssessment' field. Prioritised assessments were further added to the corresponding species within the Occurrence Dataset and the ASO Species List.

The final Extinction Risk Dataset (Extinction_risk_dataset.csv) contains all compiled extinction risk assessments (5,403 assessments) for 2,806 unique species within the ASO Species List. In addition to information on taxonomy, assessment source, scale, and publication, along with the original, standardised, and prioritised assessments.

### Data Analysis

#### *Extinction Risk Assessment Overlap and Agreement*

The sources of the species- and population-level assessments were evaluated for their contribution of unique information on ASO species extinction risk. The number of both unique and shared species was compared among sources. The degree of agreement between standardised conservation statuses was calculated using the Cohen's Kappa statistic (Cohen, 1960), using *irr* (Matthias Gamer et al., 2019). Cohen's Kappa is a measure of interrater reliability, a result of K between 0 and 1 indicates a more than chance agreement, and a K between -1 and 0 indicates less than chance agreement (Cohen, 1960). Cohen's Kappa was performed using pairwise comparisons between the two largest sources of assessments, the IUCN Red List and NatureServe, at both the species and population levels. Cohen's Kappa analyses were followed by confusion matrices using the *janitor* package (Sam Firke et al., 2024) to compare categories which are contributing to agreement or disagreement.

#### *Extinction Risk Assessment Coverage*

Examining the taxonomic, temporal, habitat, and geographic coverage of extinction risk assessments allows for the identification of potential biases in assessment effort and gaps in conservation status knowledge. Taxonomic coverage was quantified by the number of species assessed per major taxonomic group and taxonomic sub-groups. Habitat coverage was examined by comparing the number of species classified within terrestrial, marine, and inland-water habitats. It is important to examine habitat coverage as different habitats may face very distinct threats. Species with multiple habitat classifications, e.g. are classified as both marine and terrestrial species, were included in both analyses for both habitat types. Temporal patterns in assessments were examined by quantifying the accumulation of assessments over time. Assessments extracted from NatureServe did not include the assessment date, therefore NatureServe assessments were excluded from temporal analysis.

Geographic coverage of assessed species within the ASO region was evaluated to identify potential spatial biases in assessment coverage. The ASO region was rasterised at a 100 km resolution in South Pole Lambert Azimuthal Equal Area projection (ESRI:102020) and overlaid with a 10 km resolution coastline map from *rnaturalearth* (Massicotte et al., 2023). This was implemented using the *sf* (Pebesma et al., 2024) and *raster* (Murrell, 2019) packages. Assessment density was calculated using the Occurrence Dataset and summing the number of assessed species per cell. Patterns of assessments have been shown to be correlated with local species richness therefore assessment density was compared to species richness per cell, with the resulting rasters tested for correlation using the Spearman Rank Correlation Coefficient. The Spearman Rank Correlation Coefficient shows the strength and direction of a monotonic relationship, with values ranging between -1 (strong negative monotonic relationship), and 1 (strong positive monotonic relationship). Assumptions of monotonic relationships were checked and satisfied. Assessment effort was also examined looking at the proportion of assessed species to the total species richness within each cell.

*Threatened Species*

Understanding the taxonomic, habitat, and geographic coverage of threatened species is essential for targeted conservation planning. To examine the taxonomic coverage of threatened species, the proportion of species within each standardised threat category was calculated by major taxonomic groups (i.e. vertebrates and invertebrates) and taxonomic sub-groups (i.e. fishes and annelids). To investigate if there were relationships between assigned threat status and both major taxonomic groups and taxonomic sub-groups, chi-squared analysis and standardised residuals were used. These results were used to identify which taxonomic groups were more or less likely to be assigned a threat status than expected by chance. Note that species categorised as 'Extinct' and those within gymnosperm and arachnid taxon groups were excluded from analyses due to low numbers of species (<10).

The categorisation of species as Data Deficient introduces uncertainty into the analysis, as their true level of extinction risk remains unknown. To account for this uncertainty, methods established in the IUCN guidelines for reporting on the proportion of Threatened species (version 1.2) were used to estimate the lower bound, mid-point, and upper bounds of Threatened species (IUCN, 2022). The lower bound is calculated as T/*N*, where T is the number of species in the Threatened category, and *N* correlates to the number of species assessed. This assumes that none of the Data Deficient species are Threatened and is likely to underestimate the true extinction risk of a taxa. The mid-point, considered most realistic estimation of a taxa's true extinction risk (IUCN, 2022), is calculated as (T/ [*N* - EX - DD]), where EX and DD are the number of species the corresponding Extinct and Data Deficient categories. The upper bound is calculated as ([T + DD]/ [*N* - EX]) and assumes that all Data Deficient species are Threatened, which likely overestimates the true extinction risk of taxa as not all data deficient species is threatened. Range is the reported is the upper and lower bound.

The ASO mid-points for each taxon were compared to global proportions as reported by the IUCN Red List to identify taxonomic groups which may be more or less Threatened. When available, mid-points were taken from IUCN Summary Statistics, 'Table 1a: Number of species evaluated in relation to the overall number of described species, and numbers of threatened species by major groups of organisms'(IUCN, 2025b). For taxonomic groups which differed to the groupings used in the IUCN Red List Summary statistics (e.g. Fungi), or those which did not have a published midpoint in IUCN (2025) Table 1a (e.g. insects), mid-points were calculated using data from summary statistics 'Table 3: number of species in each IUCN Red List Category by the kingdom and classes downloaded from the IUCN Red List on the 18$^{th}$ of June 2025 (IUCN, 2025b).

The geographic coverage of species classified in the standardised category of Threatened within the ASO region was examined to identify areas of potential elevated threat or conservation value. Using the Occurrence Dataset, the number of Threatened species per cell were summed at a 100 km resolution as previously done for assessed species. To further account for regional variation in species richness and assessment effort, weighted species richness was calculated. Weighted species richness examines the relationship between Threatened species relative to the total richness of Data Sufficient species (Data Sufficient species = Number of species assessed - Data Deficient species) with the standardised residuals for each cell calculated. Analysis of standardised residuals enables identification of areas with more threatened species than predicted based on species richness and assessment effort (Hoffmann et al., 2010) which can be used to identify regions of high conservation value.

*Threats to Species*

For assessments published by the IUCN Red List, threat information is required for any species-level assessment classified within the Threatened categories (Critically Endangered, Endangered, and Vulnerable) as well as the Extinct categories (Extinct and Extinct in the Wild) (IUCN, 2013) Threat information is coded according to the IUCN threat classification scheme version 3.3 (IUCN, 2025a). Species with available threat data ($n$ = 775) were extracted from the IUCN Red List. The proportion of species which are affected by each threat was calculated across both major taxonomic groups and taxonomic subgroups. The proportion of species affected by each threat was calculated for each taxonomic group and taxonomic subgroup across all species assessed by the IUCN Red List globally. All assessments with available threat data were downloaded from the IUCN Red List on the 6 May 2025 (IUCN, 2025a). Proportions of species listed under each threat was calculated and compared to the ASO.

*Protected Area Coverage of Threatened Species*

Protected areas are one of the most effective strategies in aiding global biodiversity goals and protecting Threatened species if done strategically (Bolam et al., 2021; Cardillo et al., 2023; Rodrigues et al., 2004; Venter et al., 2014). A reduction of threat status over time or a lower proportion of Threatened species can be used as an indicator of effective protected-area designation and conservation management (Mace et al., 2018; Ward et al., 2024).

The relationship between protected area coverage and species conservation status was assessed by examining the extent of terrestrial and marine area-based protection afforded to species. Terrestrial protected areas included 76 Antarctic Specially Protected Areas (ASPAs) and six Antarctic Specially Managed Areas (ASMAs) (Antarctic Treaty Secretariat, 2025). Marine protected areas included two Marine Protected Areas (MPAs) established by the Commission for the Conservation of Antarctic Marine Living Resources (CAMMLR). Additionally, there are five United Nations Educational, Scientific and Cultural Organization (UNESCO) World Heritage Sites located within the ASO which encompass both marine and terrestrial environments. Shapefiles of all ASPAs and ASMAs were obtained from the Secretariat of the Antarctic Treaty Antarctic protected Areas database on the 5th of August (Antarctic Treaty Secretariat, 2025). The shapefiles of additional protected areas within the ASO including the MPAs and UNESCO World Heritage sited were downloaded through the Protected Planet database on the 1st of July 2025 (UNEP-WCMC and IUCN, 2025).

Polygons of protected areas were used to calculate the area of overlap with each species' extent of occurrence (EOO), based on records in the Occurrence Dataset. For species with at least three occurrences ($n$ = 11,845 species), their EOO was calculated following IUCN guidelines (IUCN Standards and Petitions Committee, 2024) using convex hulls in ESRI:102020 projected coordinated system. To account for spatial uncertainty in the geographic coordinate accuracy of occurrences, each protected area was given a 1 km buffer. The area and proportion of each species' EOO overlapping with any protected area in the ASO was then calculated. Differences in marine and terrestrial species protected area overlap (dependent variable) among conservation statuses (independent variable) was assessed using a Kruskal-Wallis non-parametric test (Kruskal & Wallis, 1952). Non-normality was confirmed with a Shapiro-Wilk normality test. Post-hoc pairwise comparisons were then performed using Dunn’s test (Dunn, 1964) with Benjamini-Hochberg correction.

**Appendix S2:** National and supra-national publications of population-level extinction risk assessments for species within the Antarctic Southern Ocean downloaded through the National Red List Database (ZSL and IUCN National Red List Working Group, 2022).

| Publication Name | Publishing Country or Region | Number of assessments |
|---|---|---|
| Canadian Wildlife Species at Risk | Canada | 28 |
| Carpathian List of Endangered Species | Carpathian (Central and east European) | 1 |
| Categorización de Especies Amenazadas de Fauna Silvestre | Peru | 2 |
| Červený seznam ohrožených druhů České republiky, Obratlovci | Czech Republic | 1 |
| COSEWIC- Committee on the Status of Endangered Wildlife in Canada (1989) | Canada | 2 |
| COSEWIC- Committee on the Status of Endangered Wildlife in Canada (1990) | Canada | 8 |
| COSEWIC- Committee on the Status of Endangered Wildlife in Canada (1991) | Canada | 2 |
| COSEWIC- Committee on the Status of Endangered Wildlife in Canada (1993) | Canada | 6 |
| COSEWIC- Committee on the Status of Endangered Wildlife in Canada (1994) | Canada | 2 |
| COSEWIC- Committee on the Status of Endangered Wildlife in Canada (2000) | Canada | 2 |
| COSEWIC- Committee on the Status of Endangered Wildlife in Canada (2001) | Canada | 8 |
| COSEWIC- Committee on the Status of Endangered Wildlife in Canada (2002) | Canada | 14 |
| COSEWIC- Committee on the Status of Endangered Wildlife in Canada (2003) | Canada | 10 |
| COSEWIC- Committee on the Status of Endangered Wildlife in Canada (2004) | Canada | 4 |
| COSEWIC- Committee on the Status of Endangered Wildlife in Canada (2005) | Canada | 4 |
| COSEWIC- Committee on the Status of Endangered Wildlife in Canada (2006) | Canada | 6 |
| COSEWIC- Committee on the Status of Endangered Wildlife in Canada (2007) | Canada | 4 |
| COSEWIC- Committee on the Status of Endangered Wildlife in Canada (2008) | Canada | 7 |
| Crvena knjiga sisavaca Hrvatske (Red Book of Mammals of Croatia) | Croatia | 2 |
| Eesti punane raamat | Estonia | 3 |
| Especies Amenazadas de Chile: Protejámoslas y evitemos su extinción | Chile | 1 |
| Ireland Red List No. 3: Terrestrial Mammals | Ireland | 3 |
| Korean Red List of Threatened Species | Korea | 1 |

| | | |
|---|---|---|
| La Liste rouge des espèces menacées en France 2009 | France | 16 |
| Libro Rojo de la Fauna Venezolana | Venezuela | 3 |
| Libro Rojo de la Fauna Venezolana. Tercera Edición | Venezuela | 11 |
| Libro Rojo de los Mamíferos Amenazados de la Argentina | Argentina | 57 |
| Libro Rojo de los mamiferos de Colombia | Colombia | 7 |
| Libro Rojo de los Mamíferos del Ecuador | Ecuador | 9 |
| Lista de especies ameacadas 2014 | Brazil | 11 |
| Lista de Especies en Peligro de Extinción, Amenazadas o Protegidas de la República Dominicana | The Dominican Republic | 1 |
| Lista Rossa Dei Vertebrati Italiani | Italy | 9 |
| Lista Vermelha de especies de Angola: Extintas, ameaçadas de extinção, vulneráveis e invasoras | Angola | 5 |
| Listës Së Kuqe Të Florës Dhe Faunës Së Egër, Shqipëri (Red List of Wild Flora and Fauna, Albania) | Albania | 6 |
| Mongolian Red List of Mammals | Mongolia | 2 |
| Nationally Threatened Species for Uganda | Uganda | 2 |
| New Zealand Threat Classification System lists - 2005 | New Zealand | 40 |
| Norsk Rødliste 2006 | Norway | 4 |
| Norsk rødliste for arter 2010 | Norway | 5 |
| Red Data Book of the Mammals of South Africa: A Conservation Assessment | South Africa | 31 |
| Red Data Book of the Republic of Bulgaria | Republic of Bulgaria | 1 |
| Red List of Bangladesh Volume 2: Mammals | Bangladesh | 7 |
| Red List of China's Vertebrates | China | 24 |
| Rödlistade arter i Sverige 2005 | Sweeden | 3 |
| Rödlistade arter i Sverige 2010 | Sweeden | 3 |
| Rote Liste gefährdeter Tiere Deutschlands | Germany | 4 |
| Rote Listen gefährdeter Tiere österreichs. Checklisten, Gefährdungsanalysen, Handlungsbedarf. Teil 1: Säugetiere, Vögel, Heuschrecken, Wasserkäfer, Netzflügler, Schnabelfliegen, Tagfalter | Austria | 6 |
| Status of South Asian Non-volant Small Mammals: Conservation Assessment and Management Plan (C.A.M.P.) Workshop Report | South Asia | 14 |
| Suomen lajien uhanalaisuus 2000 | Finland | 2 |
| Suomen lajien uhanalaisuus 2010 | Finland | 4 |

| | | |
|---|---|---|
| The 2016 Red List of Mammals of South Africa, Swaziland and Lesotho | South Africa, Swaziland and Lesotho | 36 |
| The 4th Japanese Red Data Book | Japan | 2 |
| The National Red List 2012 of Sri Lanka: Conservation Status of the Fauna and Flora | Sri Lanka | 19 |
| The Red Book: vertebrates in Israel | Israel | 4 |
| The Status of Nepal's Mammals: The National Red List Series | Nepal | 3 |
| Threatened species under the EPBC Act | Australia | 8 |
| UAE National Red List of Birds | United Arab Emirates | 31 |
| UAE National Red List of Herpetofauna: Amphibians & Terrestrial Reptiles, Sea Snakes & Marine Turtles | United Arab Emirates | 5 |
| UAE National Red List of Mammals: Marine and Terrestrial | United Arab Emirates | 10 |
| UAE National Red List of Marine Species: Reef-building corals, cartilaginous fishes and select bony fishes | United Arab Emirates | 23 |
| UAE National Red List of Vascular Plants | United Arab Emirates | 24 |

**<u>Appendix S3:</u>** Lexicon standardisation of different sources of extinction risk assessments to five conservation statuses (Extinct, Threatened, Not Threatened, Not Assessed, Data Deficient) where the standardised extinction risk category differed from the original category.

| Standardised Category | Original Assessment Category | Assessment Source/Publication |
|---|---|---|
| **Extinct** | Extinct, disparue ex | COSEWIC- Committee on the Status of Endangered Wildlife in Canada |
| | Extinct or missing (ausgestorben oder verschol len) | Rote Liste gefährdeter Tiere Deutschlands |
| | Extinct in the wild | IUCN Red List |
| | GX, NX, NXB | NatureServe |
| **Threatened** | Regionally Extinct; Critically Endangered; Endangered; Vulnerable | IUCN Red List |
| | Regionally Extinct; Vulnerable | La Liste rouge des espèces menacées en France |
| | Regionally Extinct; Vulnerable | Norsk Rødliste |
| | Regionally Extinct; Vulnerable | Libro Rojo de la Fauna Venezolana. Tercera Edición |
| | Regionally Extinct; Endangered; Vulnerable | Rödlistade arter i Sverige |
| | Regionally Extinct | Suomen lajien uhanalaisuus 2000 |
| | Critically Endangered; Endangered; Vulnerable | Canadian Wildlife Species at Risk |
| | Critically Endangered; Endangered; Vulnerable | Lista de especies ameacadas |
| | Critically Endangered; Endangered; Vulnerable | Red List of China's Vertebrates |
| | Critically Endangered; Endangered; Vulnerable | The 2016 Red List of Mammals of South Africa, Swaziland and Lesotho |
| | Critically Endangered; Endangered; Vulnerable | UAE National Red List of Birds |
| | Critically Endangered; Vulnerable | Rote Listen gefährdeter Tiere österreichs. Checklisten, Gefährdungsanalysen, Handlu ngsbedarf. Teil 1: Säugetiere, Vögel, Heuschrecken, Wasserkäfer, Netzflügler, Schna belfliegen, Tagfalter |

| | |
|---|---|
| Endangered; Vulnerable | Libro Rojo de los Mamíferos Amenazados de la Argentina |
| Endangered; Vulnerable | Libro Rojo de los mamiferos de Colombia |
| Endangered; Vulnerable | Libro Rojo de los Mamíferos del Ecuador |
| Endangered; Vulnerable | Lista Rossa Dei Vertebrati Italiani |
| Endangered; Vulnerable | Red Data Book of the Mammals of South Africa: A Conservation Assessment |
| Endangered; Vulnerable | The National Red List 2012 of Sri Lanka: Conservation Status of the Fauna and Flora |
| Endangered; Vulnerable | Threatened species under the EPBC Act |
| Endangered; Vulnerable | UAE National Red List of Marine Species: Reef-building corals, cartilaginous fishes and select bony fishes |
| Endangered (en peligro); Vulnerable | Libro Rojo de la Fauna Venezolana |
| Endangered | Categorización de Especies Amenazadas de Fauna Silvestre |
| Endangered | Crvena knjiga sisavaca Hrvatske (Red Book of Mammals of Croatia) |
| Endangered (gefährdet) | Rote Liste gefährdeter Tiere Deutschlands |
| Vulnerable | Especies Amenazadas de Chile: Protejámoslas y evitemos su extinción |
| Vulnerable | Ireland Red List No. 3: Terrestrial Mammals |
| Vulnerable | Korean Red List of Threatened Species |
| Vulnerable | Lista de Especies en Peligro de Extinción, Amenazadas o Protegidas de la República Dominicana |
| Vulnerable | Listës Së Kuqe Të Florës Dhe Faunës Së Egër, Shqipëri (Red List of Wild Flora and Fauna, Albania) |
| Vulnerable | Mongolian Red List of Mammals |
| Vulnerable | Red Data Book of the Republic of Bulgaria |
| Vulnerable | UAE National Red List of Herpetofauna: Amphibians & Terrestrial Reptiles, Sea Snakes & Marine Turtles |
| Vulnerable (vulnerável) vul | Lista Vermelha de especies de Angola: Extintas, ameaçadas de extinção, vulneráveis e invasoras |
| Vulnerable; Threatened local population | The 4th Japanese Red Data Book |
| Appendix I | CITES |

| | | |
|---|---|---|
| | GH; G1; G2; G3; NH; N1; N2; N3; NHB; N1B; N2B; N3B; NHN; N1N; N2N; N3N; NHM; N1M; N2M; N3M | NatureServe |
| | Endangered en voie de disparition; Threatened, menacée | COSEWIC- Committee on the Status of Endangered Wildlife in Canada |
| | Nationally Critical; Nationally Endangered; Range Restricted | New Zealand Threat Classification System |
| **Not Threatened** | Least Concern; Lower Risk/Least Concern; Lower Risk/Conservation Dependent; Lower Risk/Near Threatened; Near Threatened | IUCN Red List |
| | Lower Risk/Near Threatened; Lower Risk/Conservation Dependent | Listës Së Kuqe Të Florës Dhe Faunës Së Egër, Shqipëri (Red List of Wild Flora and Fauna, Albania) |
| | Least Concern; Near Threatened | La Liste rouge des espèces menacées en France |
| | Least Concern; Near Threatened | Red List of Bangladesh Volume 2: Mammals |
| | Least Concern; Near Threatened | Rote Listen gefährdeter Tiere österreichs. Checklisten, Gefährdungsanalysen, Handlu ngsbedarf. Teil 1: Säugetiere, Vögel, Heuschrecken, Wasserkäfer, Netzflügler, Schna belfliegen, Tagfalter |
| | Least Concern; Near Threatened | The 2016 Red List of Mammals of South Africa, Swaziland and Lesotho |
| | Least Concern; Near Threatened | UAE National Red List of Birds |
| | Least Concern; Near Threatened | UAE National Red List of Marine Species: Reef-building corals, cartilaginous fishes and select bony fishes |
| | Least Concern | Canadian Wildlife Species at Risk |
| | Least Concern | Eesti punane raamat |
| | Least Concern | Ireland Red List No. 3: Terrestrial Mammals |
| | Least Concern | Libro Rojo de la Fauna Venezolana. Tercera Edición |
| | Least Concern | Nationally Threatened Species for Uganda |
| | Least Concern | Red List of China's Vertebrates |

| | | |
|---|---|---|
| | Least Concern | Status of South Asian Non-volant Small Mammals: Conservation Assessment and Management Plan (C.A.M.P.) Workshop Report |
| | Least Concern | The Red Book: vertebrates in Israel |
| | Least Concern | The Status of Nepal's Mammals: The National Red List Series |
| | Least Concern | UAE National Red List of Vascular Plants |
| | Least Concern; RBDC; RBPM | Libro Rojo de los Mamíferos Amenazados de la Argentina |
| | Near Threatened | Lista de especies ameacadas 2014 |
| | Near Threatened | Lista Rossa Dei Vertebrati Italiani |
| | Near Threatened | Mongolian Red List of +C75Mammals |
| | Near Threatened | Norsk Rødliste |
| | Near Threatened | Red Data Book of the Mammals of South Africa: A Conservation Assessmen |
| | Near Threatened | Suomen lajien uhanalaisuus 2000 |
| | Not at risk, non en péril; Special concern, préoccupante | COSEWIC- Committee on the Status of Endangered Wildlife in Canada |
| | G4; G5; N4; N5; N4B; N5B; N4N; N5N; N4M; N5M | NatureServe |
| | Vagrant; Migrant | New Zealand Threat Classification System lists |
| **Not Assessed** | Not Evaluated; Not Applicable | IUCN Red List |
| | Not Applicable | Ireland Red List No. 3: Terrestrial Mammals |
| | Not Applicable | La Liste rouge des espèces menacées en France |
| | Not Applicable | Lista Rossa Dei Vertebrati Italiani |
| | Not Evaluated | Red Data Book of the Mammals of South Africa: A Conservation Assessmen |
| | Not Evaluated | Rote Listen gefährdeter Tiere österreichs. Checklisten, Gefährdungsanalysen, Handlungsbedarf. Teil 1: Säugetiere, Vögel, Heuschrecken, Wasserkäfer, Netzflügler, Schnabelfliegen, Tagfalter |
| | Not Evaluated | Status of South Asian Non-volant Small Mammals: Conservation Assessment and Management Plan (C.A.M.P.) Workshop Report |
| | Not Evaluated | The 2016 Red List of Mammals of South Africa, Swaziland and Lesotho |

| | | |
|---|---|---|
| | GNR; GNA; NNR; NNA; NNRB; NNAB; NNRN; NNAN; NNRM; NNAM | NatureServe |
| **Data Deficient** | Data Deficient, données insuffisantes | COSEWIC- Committee on the Status of Endangered Wildlife in Canada |
| | Data Deficient (datenlage unklar) | Rote Liste gefährdeter Tiere Deutschlands |
| | GU; NU; NUB; NUN; NUM | NatureServe |

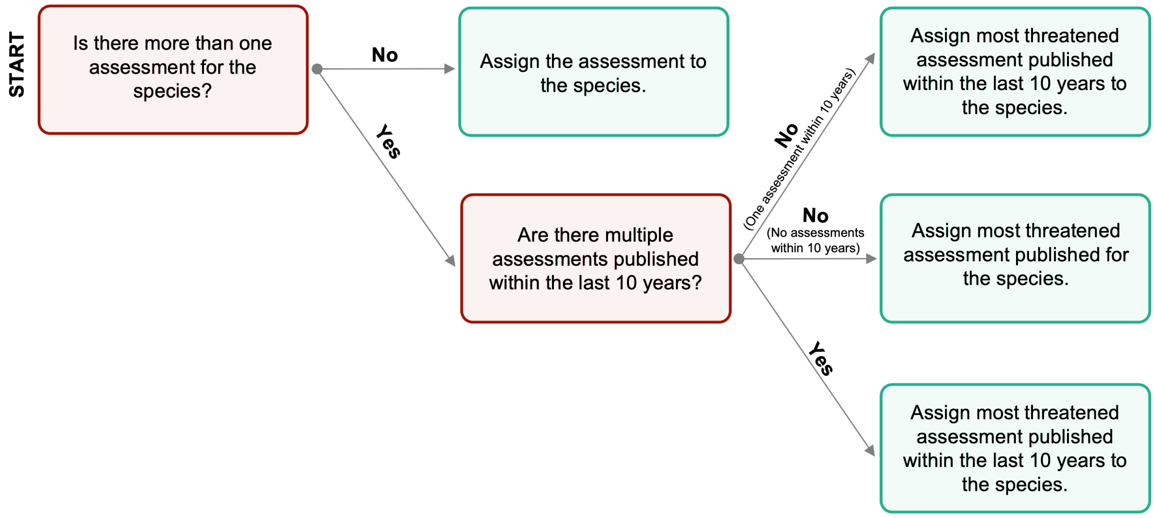


**<u>Appendix S4:</u>** Workflow used to prioritise Antarctic and Southern Ocean species extinction risk conservation statuses where multiple assessments for a species was present.

**Appendix S5:** Fields present within the Occurrence Dataset, Antarctic and Southern Ocean (ASO) Species List, and the Extinction Risk Datasets. An asterisk (*) indicates fields that align with Darwin Core Standards (Wieczorek et al., 2012).

| Field | Description | Dataset/s |
|---|---|---|
| scientificName* | The binomial scientific name at the species level according to the taxonomic source in 'nameAccordingTo'. | All |
| originalNameUsage* | The taxon name, with authorship and date information if known, as it originally appeared as per the parent dataset in 'datasetName'. | Occurrence Dataset, ASO Species List |
| kingdom*/ phylum* / class*/ order*/ family*/ genus* | Higher level taxonomy for the 'scientificName' according to the taxonomic source in 'nameAccordingTo'. | All |
| nameAccordingTo* | The reference to the source for which the taxon identification has come from. Terms: GBIF Backbone Taxonomy; Not Corrected. | All |
| taxonRank* | The taxonomic rank of the most specific name identified in 'scientificName'. | Occurrence Dataset, ASO Species List |
| occurrenceID* | Unique identifier for the occurrence verbatim as per the parent dataset in 'datasetName'. | Occurrence Dataset |
| occurrenceStatus* | A statement about the presence or absence of a taxa at a location as per the parent dataset in 'datasetName'. Terms: Present; Absent. | Occurrence Dataset |
| datasetKey* | The DOI or specific identifier for the download of the occurrence records as per the parent dataset in 'datasetName'. | Occurrence Dataset, ASO Species List |
| datasetName* | The name identifying the dataset or repository which the record was derived. Terms: GBIF; OBIS; SIVFLORA; Biodiversity of ice-free Antarctica; TerrANTALife. | Occurrence Dataset, ASO Species List |
| taxonID* | Identifier for species verbatim as in TerrANTALife. | All |
| eventDate* | The date when the occurrence was observed, as per the parent dataset in 'datasetName'. Only years have been retained in 'yyyy' format. | Occurrence Dataset |
| decimalLatitude* | The geographic latitude in decimal degrees as per the parent dataset in 'datasetName'. | Occurrence Dataset |
| decimalLongitude* | The geographic longitude in decimal degrees as per the parent dataset in 'datasetName'. | Occurrence Dataset |

| | | |
|---|---|---|
| geodeticDatum* | The coordinate reference system upon which the geographic coordinates are given in 'decimalLatitude' and 'decimalLongitude' as per the parent dataset in 'datasetName'. | Occurrence Dataset |
| coordinateUncertaintyInMeters* | The horizontal distance in meters from the given 'decimalLatitude' and 'decimalLongitude' describing the smallest circle containing the whole of the occurrence location as per the parent dataset in 'datasetName'. | Occurrence Dataset |
| habitat* | A category or description of the habitat in which the occurrence or taxon occurred. Terms: Marine; Terrestrial; Inland-water and their combinations. | All |
| higherGeographyID* | An identifier for the geographic region within which the occurrence/s was recorded. Terms: Antarctic (south of 60°S); sub-Antarctic and Southern Ocean (north of 60°S) and their combinations. | Occurrence Dataset |
| basisOfRecord* | The specific nature of the record as per the parent dataset in 'datasetName'. Terms: HumanObservation; MachineObservation; PreservedSpecimen; MaterialSample; MaterialCitation; LivingSpecimen; Occurrence; Observation. | Occurrence Dataset |
| taxonCategory | Taxonomic sub-Groups assigned to species. Terms: Mammals; Birds; Fishes; Reptiles; Corals; Crustaceans; Molluscs; Echinoderms; Insects; Sponges; Annelids; Arachnids; Ferns and Allies; Flowering Plants; Mosses; Gymnosperms; Fungi. | All |
| standardisedAssessment | Standardised terminology of 'verbatimAssessment'. Terms: Extinct; Threatened; Not Threatened; Data Deficient; Not Assessed. | Extinction Risk Dataset |
| selectedAssessment | The prioritised 'standardisedAssessment' assigned to a species. Terms: Extinct; Threatened; Not Threatened; Data Deficient; Not Assessed. | All |
| verbatimAssessmentScope | Assessment scope verbatim as per the 'assessmentSource'. | Extinction Risk Dataset |
| assessmentYear | Year assessment was conducted as per the 'assessmentSource' in yyyy format. | Extinction Risk Dataset |
| assessmentSystem | The system that was used to assign an extinction risk category. Fields: IUCN; non-IUCN; Modified IUCN. | Extinction Risk Dataset |
| assessmentSource | The database where the original extinction risk assessment was downloaded from. Terms: IUCN Red List of Threatened Species; National Red List Database; NatureServe; Literature. | Extinction Risk Dataset |

| | | |
|---|---|---|
| assessmentScope | The geographic scale of the assessment region. Fields: Species-level (Global); Population-level (sub-Global). | Extinction Risk Dataset |
| assessmentPublication | The original assessment publication title that the extinction risk was derived from. This is additional, more specific information to accompany the ‘assessmentSource’. | Extinction Risk Dataset |
| verbatimAssessmentScope | The original scale of the assessment verbatim as per the ‘assessmentSource’. | Extinction Risk Dataset |
| gbifID | Alpha-numeric identifier from GBIF download. | Occurrence Dataset |
| Locality* | Description of the place of occurrence. | Occurrence Dataset |

**Appendix S6:** Current protected areas found either partly or wholly within the Antarctic and Southern Ocean Marine Ecosystem Assessment of the Southern Ocean (MEASO) region. Antarctic Specially Managed Areas and Antarctic Specially Protected Areas were sourced from Antarctic Treaty Secretariat (2025) protected areas database, and all additional protected areas from Protected Planet website (UNEP-WCMC and IUCN, 2025).

| Protected Area | Type | Habitat |
|---|---|---|
| ASMA 1: Admiralty Bay, King George Island | Antarctic Specially Managed Area | Terrestrial |
| ASMA 2: McMurdo Dry Valleys, Southern Victoria Land | Antarctic Specially Managed Area | Terrestrial |
| ASMA 3: Deception Island | Antarctic Specially Managed Area | Terrestrial |
| ASMA 4: Amundsen-Scott South Pole Station, South Pole | Antarctic Specially Managed Area | Terrestrial |
| ASMA 5: Larsemann Hills, East Antarctica | Antarctic Specially Managed Area | Terrestrial |
| ASMA 7: Southwest Anvers Island and Palmer Basin | Antarctic Specially Managed Area | Terrestrial |
| ASPA 101: Taylor Rookery, Mac. Robertson Land | Antarctic Specially Protected Area | Terrestrial |
| ASPA 102: Rookery Islands, Holme Bay, Mac. Robertson Land | Antarctic Specially Protected Area | Terrestrial |
| ASPA 103: Ardery Island and Odbert Island, Budd Coast, Wilkes Land, East Antarctica | Antarctic Specially Protected Area | Terrestrial |
| ASPA 104: Sabrina Island, Balleny Islands | Antarctic Specially Protected Area | Terrestrial |
| ASPA 105: Beaufort Island, McMurdo Sound, Ross Sea | Antarctic Specially Protected Area | Terrestrial |
| ASPA 106: Cape Hallett, Northern Victoria Land, Ross Sea | Antarctic Specially Protected Area | Terrestrial |
| ASPA 107: Emperor Island, Dion Islands, Marguerite Bay, Antarctic Peninsula | Antarctic Specially Protected Area | Terrestrial |
| ASPA 108: Green Island, Berthelot Islands, Antarctic Peninsula | Antarctic Specially Protected Area | Terrestrial |
| ASPA 109: Moe Island, South Orkney Islands | Antarctic Specially Protected Area | Terrestrial |
| ASPA 110: Lynch Island, South Orkney Islands | Antarctic Specially Protected Area | Terrestrial |
| ASPA 111: Southern Powell Island and adjacent islands, South Orkney Islands | Antarctic Specially Protected Area | Terrestrial |
| ASPA 112: Coppermine Peninsula, Robert Island, South Shetland Islands | Antarctic Specially Protected Area | Terrestrial |
| ASPA 113: Litchfield Island, Arthur Harbor, Anvers Island, Palmer Archipelago | Antarctic Specially Protected Area | Terrestrial |
| ASPA 115: Lagotellerie Island, Marguerite Bay, Graham Land | Antarctic Specially Protected Area | Terrestrial |
| ASPA 116: New College Valley, Caughley Beach, Cape Bird, Ross Island | Antarctic Specially Protected Area | Terrestrial |
| ASPA 117: Avian Island, Marguerite Bay, Antarctic Peninsula | Antarctic Specially Protected Area | Terrestrial |
| ASPA 119: Davis Valley and Forlidas Pond, Dufek Massif, Pensacola Mountains | Antarctic Specially Protected Area | Terrestrial |
| ASPA 120: Pointe-Géologie Archipelago, Terre Adélie | Antarctic Specially Protected Area | Terrestrial |

| | | |
|---|---|---|
| ASPA 121: Cape Royds, Ross Island | Antarctic Specially Protected Area | Terrestrial |
| ASPA 122: Arrival Heights, Hut Point Peninsula, Ross Island | Antarctic Specially Protected Area | Terrestrial |
| ASPA 123: Barwick and Balham Valleys, Southern Victoria Land | Antarctic Specially Protected Area | Terrestrial |
| ASPA 124: Cape Crozier, Ross Island | Antarctic Specially Protected Area | Terrestrial |
| ASPA 125: Fildes Peninsula, King George Island (25 de Mayo) | Antarctic Specially Protected Area | Terrestrial |
| ASPA 126: Byers Peninsula, Livingston Island, South Shetland Islands | Antarctic Specially Protected Area | Terrestrial |
| ASPA 127: Haswell Island | Antarctic Specially Protected Area | Terrestrial |
| ASPA 128: Western shore of Admiralty Bay, King George Island, South Shetland Islands | Antarctic Specially Protected Area | Terrestrial |
| ASPA 129: Rothera Point, Adelaide Island | Antarctic Specially Protected Area | Terrestrial |
| ASPA 131: Canada Glacier, Lake Fryxell, Taylor Valley, Victoria Land | Antarctic Specially Protected Area | Terrestrial |
| ASPA 132: Potter Peninsula, King George Island (Isla 25 de Mayo), South Shetland Islands | Antarctic Specially Protected Area | Terrestrial |
| ASPA 133: Harmony Point, Nelson Island, South Shetland Islands | Antarctic Specially Protected Area | Terrestrial |
| ASPA 134: Cierva Point and offshore islands, Danco Coast, Antarctic Peninsula | Antarctic Specially Protected Area | Terrestrial |
| ASPA 135: North-east Bailey Peninsula, Budd Coast, Wilkes Land | Antarctic Specially Protected Area | Terrestrial |
| ASPA 136: Clark Peninsula, Budd Coast, Wilkes Land, East Antarctica | Antarctic Specially Protected Area | Terrestrial |
| ASPA 137: Northwest White Island, McMurdo Sound | Antarctic Specially Protected Area | Terrestrial |
| ASPA 138: Linnaeus Terrace, Asgard Range, Victoria Land | Antarctic Specially Protected Area | Terrestrial |
| ASPA 139: Biscoe Point, Anvers Island, Palmer Archipelago | Antarctic Specially Protected Area | Terrestrial |
| ASPA 140: Parts of Deception Island, South Shetland Islands | Antarctic Specially Protected Area | Terrestrial |
| ASPA 141: Yukidori Valley, Langhovde, Lützow-Holm Bay | Antarctic Specially Protected Area | Terrestrial |
| ASPA 142: Svarthamaren | Antarctic Specially Protected Area | Terrestrial |
| ASPA 143: Marine Plain, Mule Peninsula, Vestfold Hills, Princess Elizabeth Land | Antarctic Specially Protected Area | Marine |
| ASPA 145: Port Foster, Deception Island, South Shetland Islands | Antarctic Specially Protected Area | Terrestrial |
| ASPA 146: South Bay, Doumer Island, Palmer Archipelago | Antarctic Specially Protected Area | Terrestrial |
| ASPA 147: Ablation Valley and Ganymede Heights, Alexander Island | Antarctic Specially Protected Area | Terrestrial |
| ASPA 148: Mount Flora, Hope Bay, Antarctic Peninsula | Antarctic Specially Protected Area | Terrestrial |
| ASPA 149: Cape Shirreff and San Telmo Island, Livingston Island, South Shetland Islands | Antarctic Specially Protected Area | Terrestrial |
| ASPA 150: Ardley Island, Maxwell Bay, King George Island (25 de Mayo) | Antarctic Specially Protected Area | Terrestrial |

| | | |
|---|---|---|
| ASPA 151: Lions Rump, King George Island, South Shetland Islands | Antarctic Specially Protected Area | Terrestrial |
| ASPA 154: Botany Bay, Cape Geology, Victoria Land | Antarctic Specially Protected Area | Terrestrial |
| ASPA 155: Cape Evans, Ross Island | Antarctic Specially Protected Area | Terrestrial |
| ASPA 156: Lewis Bay, Mount Erebus, Ross Island | Antarctic Specially Protected Area | Terrestrial |
| ASPA 157: Backdoor Bay, Cape Royds, Ross Island | Antarctic Specially Protected Area | Terrestrial |
| ASPA 158: Hut Point, Ross Island | Antarctic Specially Protected Area | Terrestrial |
| ASPA 159: Cape Adare, Borchgrevink Coast | Antarctic Specially Protected Area | Terrestrial |
| ASPA 160: Frazier Islands, Windmill Islands, Wilkes Land, East Antarctica | Antarctic Specially Protected Area | Terrestrial |
| ASPA 161: Terra Nova Bay, Ross Sea | Antarctic Specially Protected Area | Terrestrial |
| ASPA 162: Mawson’s Huts, Cape Denison, Commonwealth Bay, George V Land, East Antarctica | Antarctic Specially Protected Area | Terrestrial |
| ASPA 163: Dakshin Gangotri Glacier, Dronning Maud Land | Antarctic Specially Protected Area | Terrestrial |
| ASPA 164: Scullin and Murray Monoliths, Mac.Robertson Land | Antarctic Specially Protected Area | Terrestrial |
| ASPA 165: Edmonson Point, Wood Bay, Ross Sea | Antarctic Specially Protected Area | Terrestrial |
| ASPA 166: Port-Martin, Terre-Adélie | Antarctic Specially Protected Area | Terrestrial |
| ASPA 167: Hawker Island, Princess Elizabeth Land | Antarctic Specially Protected Area | Terrestrial |
| ASPA 168: Mount Harding, Grove Mountains, East Antarctica | Antarctic Specially Protected Area | Terrestrial |
| ASPA 169: Amanda Bay, Ingrid Christensen Coast, Princess Elizabeth Land, East Antarctica | Antarctic Specially Protected Area | Terrestrial |
| ASPA 170: Marion Nunataks, Charcot Island, Antarctic Peninsula | Antarctic Specially Protected Area | Terrestrial |
| ASPA 171: Narebski Point, Barton Peninsula, King George Island | Antarctic Specially Protected Area | Terrestrial |
| ASPA 172: Lower Taylor Glacier and Blood Falls, McMurdo Dry Valleys, Victoria Land | Antarctic Specially Protected Area | Terrestrial |
| ASPA 173: Cape Washington and Silverfish Bay, Terra Nova Bay, Ross Sea | Antarctic Specially Protected Area | Terrestrial |
| ASPA 174: Stornes, Larsemann Hills, Princess Elizabeth Land | Antarctic Specially Protected Area | Terrestrial |
| ASPA 175: High Altitude Geothermal sites of the Ross Sea region | Antarctic Specially Protected Area | Terrestrial |
| ASPA 176: Rosenthal Islands, Anvers Island, Palmer Archipelago | Antarctic Specially Protected Area | Terrestrial |
| ASPA 177: Léonie Islands and South-East Adelaide Island, Antarctic Peninsula | Antarctic Specially Protected Area | Terrestrial |
| ASPA 178: Inexpressible Island and Seaview Bay, Ross Sea | Antarctic Specially Protected Area | Terrestrial |
| ASPA 179: Parts of Western Sør Rondane Mountains, Dronning Maud Land, East Antarctica | Antarctic Specially Protected Area | Terrestrial |

| | | |
|---|---|---|
| ASPA 180: Danger Islands Archipelago, North-eastern Antarctic Peninsula | Antarctic Specially Protected Area | Terrestrial |
| ASPA 181: Farrier Col, Horseshoe Island, Marguerite Bay | Antarctic Specially Protected Area | Terrestrial |
| ASPA 182: Western Bransfield Strait and Eastern Dallman Bay | Antarctic Specially Protected Area | Terrestrial |
| Macquarie Island | Australia Marine Park | Marine |
| Antipodes Transect | Benthic Protection Area | Marine |
| Arrow Plateau | Benthic Protection Area | Marine |
| Bounty Heritage | Benthic Protection Area | Marine |
| Campbell East | Benthic Protection Area | Marine |
| Campbell Heritage | Benthic Protection Area | Marine |
| Sub-Antarctic Deep | Benthic Protection Area | Marine |
| Blink | Benthic protection area | Marine |
| 15JA2 b (Bollons) | Closed Seamount Area | Marine |
| 15JA2 c | Closed Seamount Area | Marine |
| 15JA2 e | Closed Seamount Area | Marine |
| 4C2 a | Closed Seamount Area | Marine |
| 4C2 e | Closed Seamount Area | Marine |
| Heard Island and McDonald Islands | Commonwealth Marine Reserve | Marine |
| Prince Edward Island Marine Protected Area | Marine Protected Area | Marine |
| South Georgia and South Sandwich Islands Marine Protected Area | Marine Protected Area | Marine |
| Ross Sea Region Marine Protected Area | Marine Protected Area (CCAMLR) | Marine |
| South Orkney Islands Southern Shelf Marine Protected Area | Marine Protected Area (CCAMLR) | Marine |
| Tristan da Cunha | Marine Protection Zone | Marine |
| Auckland Islands - Motu Maha | Marine Reserve | Marine |
| Moutere Hauriri / Bounty Islands | Marine Reserve | Marine |
| Moutere Ihupuku / Campbell Island | Marine Reserve | Marine |
| Moutere Mahue / Antipodes Island | Marine Reserve | Marine |
| Yaganes | National Marine Park | Marine |
| Terres Australes Françaises | National Nature Reserve | Marine and Terrestrial |
| Islas Diego Ramírez y Paso Drake | National Park | Terrestrial |
| Moutere Hauriri / Bounty Islands | Nature Reserve | Terrestrial |

| | | |
|---|---|---|
| Snares | Nature Reserve | Terrestrial |
| Moutere Mahue / Antipodes Island | Nature Reserve | Terrestrial |
| Bouvetøya (Antarctic) | Nature Reserve | Terrestrial |
| Réserve Naturelle Nationale des Terres Australes Française | Ramsar Site, Wetland of International Importance | Terrestrial |
| Yaganes | Strict National Marine Reserve | Marine |
| Gough and Inaccessible Islands | World Heritage Site | Marine and Terrestrial |
| French Austral Lands and Seas | World Heritage Site | Marine and Terrestrial |
| Heard and McDonald Islands | World Heritage Site | Marine and Terrestrial |
| Macquarie Island | World Heritage Site | Marine and Terrestrial |
| New Zealand Sub-Antarctic Islands | World Heritage Site | Marine and Terrestrial |

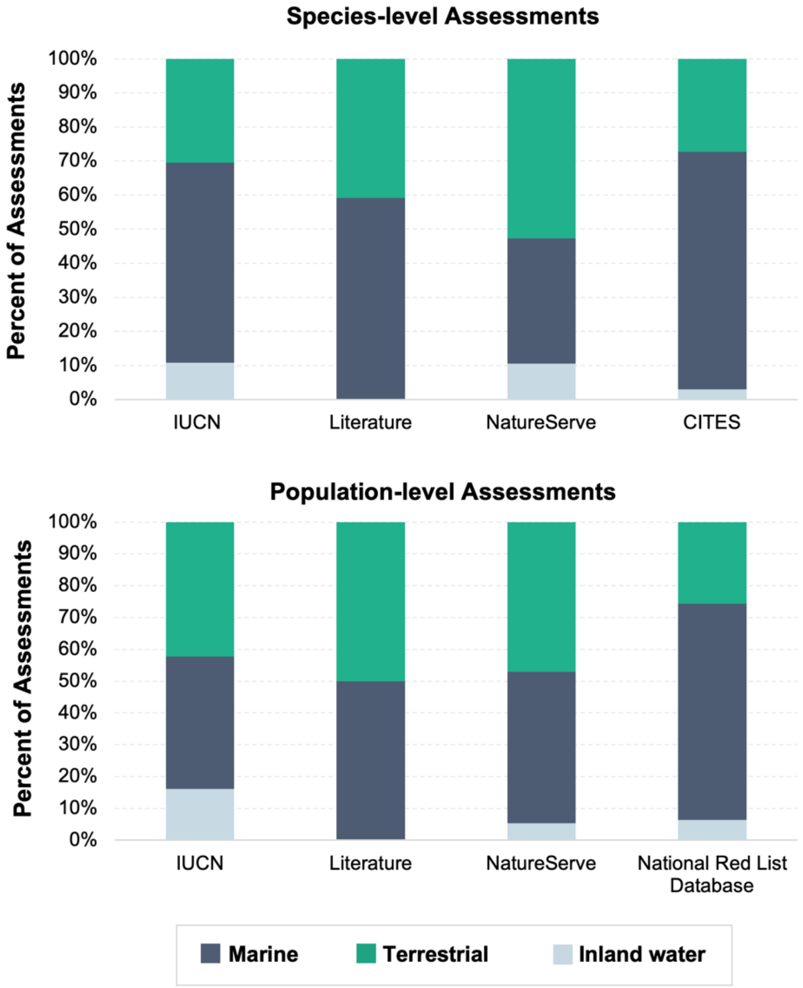


**<u>Appendix S7:</u>** Habitat coverage of extinction risk assessments by source for assessments conducted at the species- and population-levels. Note that a species may be listed under more than one habitat type.

**Appendix S8:** Confusion matrix results of extinction risk agreement between different sources and scales of extinction risk assessments.

**Species-level Assessments:** IUCN Red List and NatureServe

| | NatureServe | | | |
|---|---|---|---|---|
| **IUCN** | **Extinct** | **Threatened** | **Not Threatened** | **Data Deficient** |
| **Extinct** | 0 | 0 | 1 | 0 |
| **Threatened** | 0 | 18 | 17 | 0 |
| **Not Threatened** | 0 | 6 | 260 | 1 |
| **Data Deficient** | 0 | 0 | 12 | 0 |

**Population-level Assessments:** IUCN Red List and NatureServe

| | NatureServe | | | |
|---|---|---|---|---|
| **IUCN** | **Extinct** | **Threatened** | **Not Threatened** | **Data Deficient** |
| **Extinct** | 0 | 0 | 0 | 0 |
| **Threatened** | 0 | 4 | 6 | 5 |
| **Not Threatened** | 0 | 24 | 229 | 68 |
| **Data Deficient** | 0 | 0 | 0 | 4 |

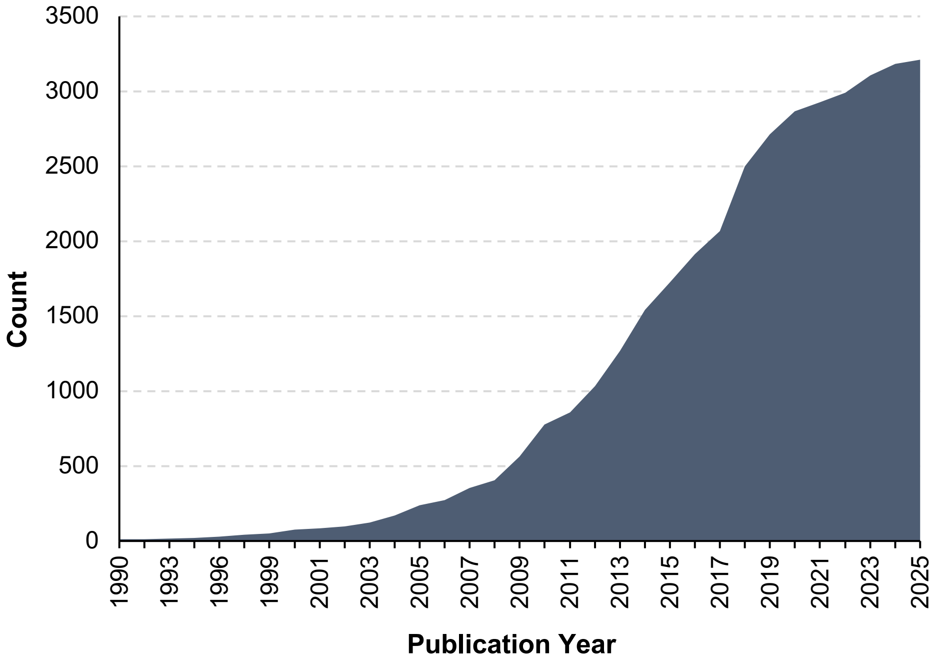


**<u>Appendix S9:</u>** Cumulative publication dates of extinction risk assessments for Antarctic and Southern Ocean Species (ASO). Extinction risk assessments were sourced from the IUCN Red list (IUCN, 2025a), CITES (CITES, 2025), the National Red List Database (ZSL and IUCN National Red List Working Group, 2022), and literature searches. Assessments for ASO species from NatureServe were excluded from this analysis due to the absence of assessment dates.

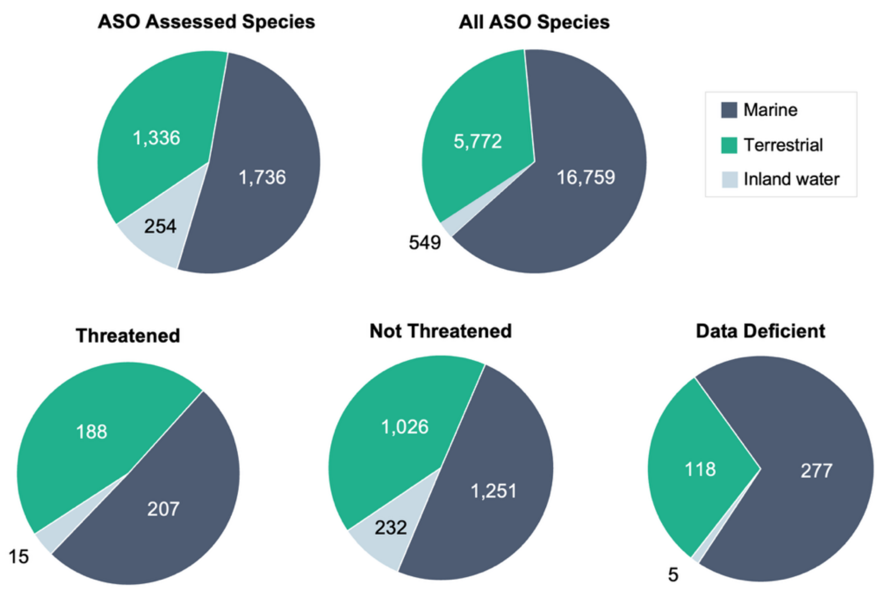


**Appendix S10:** Distribution of extinction risk assessments and conservation statuses for Antarctic and Southern Ocean (ASO) species across different habitats (marine, terrestrial and/or inland-water). These are compared the total number of ASO species within each habitat type.

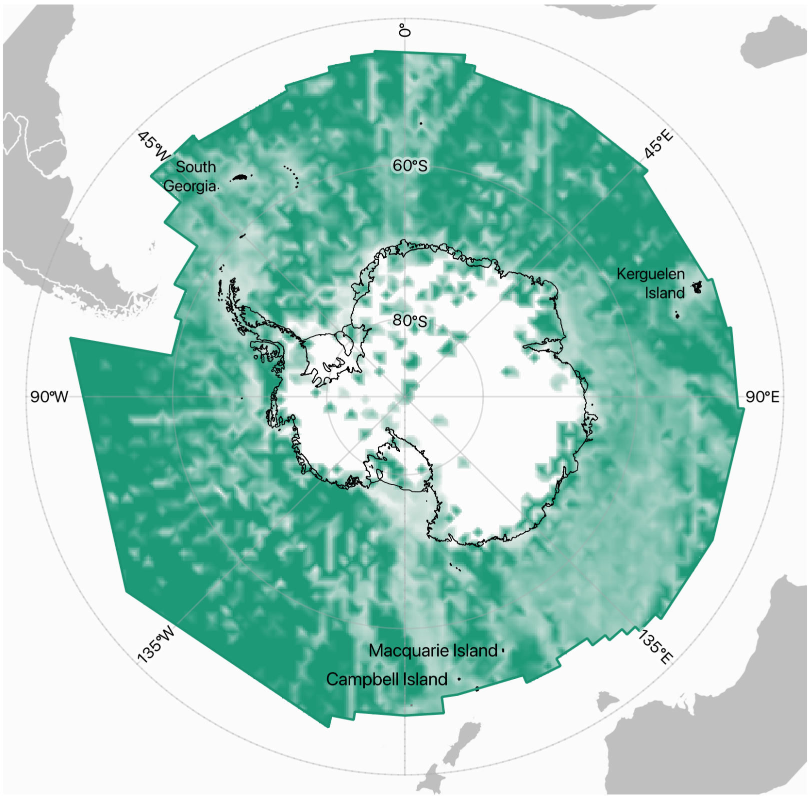


**Appendix S11:** Geographic coverage of assessment effort for Antarctic and Southern Ocean Species. Effort was measured as the number of species with assessments per 100 $km^2$ cell as a proportion of species richness

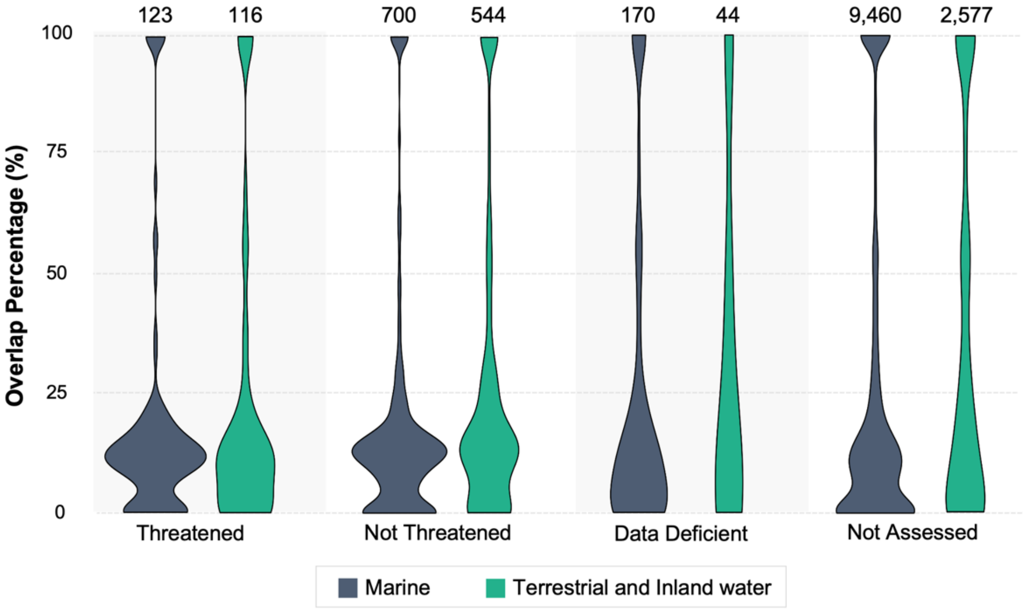


**Appendix S12:** The overlap between the extent of occurrence of species distributions (EOO) and terrestrial (including Inland-water habitats) and marine Antarctic and Southern Ocean protected areas for each standardised conservation statuses.

**<u>Appendix S13:</u>** Dunn's test *p*-value results between the overlap between the extent of occurrence of species distributions and terrestrial and marine Antarctic and Southern Ocean protected areas for each standardised threat category.

**Terrestrial**

| | Threatened | Not Threatened | Data Deficient |
|---|---|---|---|
| **Threatened** | | | |
| **Not Threatened** | 0.12 | | |
| **Data Deficient** | 0.46 | 0.80 | |
| **Not Assessed** | 0.46 | 0.03* | 0.01* |

* Indicates significant *p*-value (alpha = 0.05)

**Marine**

| | Threatened | Not Threatened | Data Deficient |
|---|---|---|---|
| **Threatened** | | | |
| **Not Threatened** | 1 | | |
| **Data Deficient** | 1 | 1 | |
| **Not Assessed** | 0.96 | 1 | 1 |
